# Effect of Ethyl Methane Sulfonate Mutagenesis on Phenological, Yield-Related and Yield Traits in Cowpea *(Vigna unguiculata* (L.) Walp)

**DOI:** 10.64898/2026.04.07.717099

**Authors:** Henry K. Mensah, Ralieva A.K. Norety, Isaac K. Asante, Felicia Oppong

**Affiliations:** Department of Plant and Environmental Biology, University of Ghana, Accra, Ghana

**Author notes:** Corresponding author (HKM). These authors contributed equally to this work (Henry K. Mensah, Ralieva A.K. Norety). These authors also contributed equally to this work (Isaac K. Asante, Felicia Oppong).

## Abstract

This study investigated the mutagenic effects of ethyl methane sulfonate (EMS) on the M□ generation in cowpea (*Vigna unguiculata* (L.) Walp.) cultivar ‘*Wang Kae’*. A total of 275 M□ seeds were treated with EMS concentrations of 20 mM, 40 mM, and 80 mM (75 seeds per treatment) by soaking for six hours, while 50 untreated seeds served as the control (0 mM). Phenological, yield-related and yield traits were recorded, and data were analysed using Jamovi 2.7.15 and JASP 0.95.4.0 through one-way ANOVA with post hoc contrast, principal component biplot, and cluster analyses. No optimal mutagenic concentration (LD50) was identified. Seed germination and seedling survival rates increased with increasing EMS concentration, ranging from 70.00% and 62.00% in the control (0 mM) to 89.33% and 74.67% at 80 mM, following the trend 0 mM < 20 mM < 40 mM < 80 mM. Significant differences (P < 0.05) were observed among treatments for all phenological traits, pod length, locule number, seed traits, and yield per plant. Yield was significantly higher (P = 0.047) at 20 mM (61.19 ± 3.34 g) compared to the control. Contrast analysis identified genotypes B33 and D56 as the most productive mutants, with yields of 125.44 g and 111.85 g, respectively. Principal component analysis extracted eighteen components, with the first four cumulatively explaining 50.60% of total variation. Biplot analysis of PC1 and PC2 captured all phenological traits, key seed traits, and yield attributes, highlighting the superior performance of B33 and D56. Cluster analysis partitioned the 190 genotypes into six groups, with B33 and D56 constituting distinct clusters. EMS mutagenesis effectively induced heritable phenotypic variation, with putative superior genotypes identified for advancement to M□ and evaluation in replicated multi-environment trials toward the development of farmer- and consumer-preferred cowpea varieties.

## Introduction

*Cowpea* (*Vigna unguiculata* [L.] Walp.) is a widely cultivated and nutritionally important legume, predominantly grown in tropical and subtropical regions, particularly in Sub-Saharan Africa, Asia, and Central and South America, while also being adapted to certain temperate zones such as the Mediterranean basin and the southern regions of the United States [1,2]. Cowpea (*Vigna unguiculata* L. Walp.), is an important staple food legume and cheap source of protein for many Africans in the low-land humid and dry savannah tropics. It is an excellent substitute for animal proteins by resource-poor people and vegetarians because of its high seed protein content (about 25%) and rich amino acids [3]. Indeed, some cultivars with seed protein content of about 30%, close to that obtained for soybean (*Glycine max*) have been reported [4,5]. Due to a decline in cowpea market value caused by pests, pathogens, and a lack of genetic variability, many scientists worldwide have sought to develop cowpea varieties that meet the taste and demand of a growing population [6–9]. Previous studies have demonstrated substantial genetic variability in cowpea for agronomic and economically important traits, including protein and mineral composition, underscoring the need to broaden the genetic base through breeding or mutagenesis [10,11]. The most noteworthy studies have used mutagens to increase genetic variability [6–9,12–14]. Ethyl Methane Sulfonate (EMS), one of the most effective mutagens, has been successfully used by several researchers in West Africa to improve genetic variability in cowpea [8,15].

A mutation is a heritable alteration in the structure or nucleotide sequence of a gene that arises suddenly and can be transmitted to subsequent generations. It results from changes in DNA such as substitutions, insertions, deletions, or rearrangements, and may occur naturally through replication or repair errors or be intentionally induced by physical and chemical mutagens. In plants, mutations can be fixed in both seed-propagated and vegetatively propagated species, allowing stable transmission of new traits. Consequently, mutations constitute a fundamental source of genetic variation and play a critical role in plant breeding and crop improvement programs [16,17]. To create functional mutants in various crop species, a variety of chemical mutagens have been used. Artificial mutation induction through chemical mutagens, including colchicine (Col), ethyl methanesulfonate (EMS) and sodium azide (SA), has been widely recognized as an effective approach for enhancing genetic variability in plants. This technique facilitates the improvement of specific traits while preserving the inherent desirable attributes of the plant, as documented by several researchers [16]. Chemical mutagens generally produce induced mutations which lead to base pair substitutions, especially GC→AT resulting in amino acid changes, which change the function of proteins but do not abolish their functions as like deletions or frame shift mutations. These chemo mutagens induce a broad variation in morphological characters when compared to normal plants [18].

Ethyl methane sulfonate is a colorless mutagenic, carcinogenic and teratogenic chemical compound with the chemical formula CH_3_OSO_2_C_2_H_5_. It is formed from the condensation of ethanal with methane sulfonic acid [19]. Ethyl methane sulphonate is an alkylating chemical mutagen that has been widely utilized in plant breeding because it can generate a high frequency of gene mutations. EMS produces O_6_-ethyl guanine by alkylating guanine residues. This matches thymine (T) but not cytosine (C) as a result, unrepaired alkylation replication occurs. A damaged pair A/T will effectively replace the G/C base pair. This mechanism foresees a strong G/C to A/T bond shortly [20]. According to [21,22], EMS is a monofunctional ethylating agent that has been reported to be mutagenic in a range of genetic test systems, ranging from viruses to mammals. Effectiveness and efficiency mean the extent to which the mutagenic agent produced a desired effect on the plant which may be positive or negative effect. [23] reported the order of effectiveness of chemical mutagens as hydrazine hydrate (HZ)> sodium azide (SA)> ethyl methane sulphonate (EMS) as they did a systematic and comparative study of induced chlorophyll mutation using HZ, SA and EMS and found that EMS treatments caused the highest frequency of chlorophyll mutations followed by HZ and then SA. [23] & [24] reported that mutations of chemical mutagens and their effectiveness is dose-dependent and increase with the concentration of mutagens. [24] reported that lower concentrations of EMS are more effective than higher concentrations. Ethyl methane sulfonate is used by plant breeders because of its ease of use, good penetration, reproducibility, high mutation frequency and lack of disposal issues.

According to [6], plant breeders’ major goal is to create crops that outperform existing cultivars in terms of production and quality, and this is dependent on the availability of genetic variation, preferably in the primary gene pool. Genetic variety is necessary for crop improvement. Plant breeders, therefore, have begun to use mutation breeding more frequently such as [13] who in their research induced variations using EMS in cowpea (*Vigna unguiculata* L. Walp Var. *Asontem*) in the M_1_ and M_2_ generations at different concentrations of EMS (0.0, 0.2, 0.4, 0.6, and 0.8%) for 16h which showed effective variations in traits such as germination and percentage survival, yield and yield related traits in M_1_ population. [25] and [26] both subjected cowpea to EMS at different does for 6 hours, also reported higher number of pods per plant, early and late phenological traits, changes in pigmentation of leaf, pod and flower colour, plant height, increased pod length and high seed weights as a result of EMS-induced mutations. During the M_1_ generation, only dominant or co-dominant mutations are easily detected by phenotypic observations [27]. Some viable mutations in the M_1_ generation include reduction in plant height, pollen sterility, late or early flowering and podding. Furthermore, most previous studies have focused on later generations (M□ and beyond), where mutations are more stably expressed, with fewer studies providing detailed analyses of variability in the M□ generation. Understanding the immediate effects of EMS in the M□ population is crucial for optimizing mutagenesis protocols and identifying early indicators of useful genetic variation.

Therefore, this study aimed to evaluate the effects of different EMS concentrations on seed germination and survival, phenological, yield-related and yield traits in the M□ generation of cowpea cultivar *’Wang Kae’*, with the objective of identifying putative superior genotypes for advancement to the M□ generation and subsequent evaluation in replicated multi-environment trials. The ultimate goal was to support the development of improved cowpea varieties with farmer- and consumer-preferred traits for integration into breeding programs. By characterizing the phenotypic variability induced at the M□ stage, this study contributes to refining EMS-based mutation breeding strategies in cowpea.

## Materials & Methods

### Experimental site, material and design

The field study was carried out at the Department of Plant and Environmental Biology, University of Ghana, from early April to July 2025. A total of 275 seeds of the ‘*Wang Kae*’ cowpea variety (M□ generation) were used in this study. Seventy-five seeds each were treated with three different concentrations of ethyl methane sulfonate (EMS): 20 mM, 40 mM, and 80 mM, and were soaked in the respective EMS solutions for six hours. Fifty seeds were soaked without treatment and served as the control (0 mM EMS). Genotypes were assigned alphanumeric labels to indicate both the EMS treatment level and the individual plant number. The alphabetic prefix represented the EMS concentration, while the numeric suffix denoted the individual plant within that treatment group. Specifically, ‘***A***’ represented the control group (0 mM EMS), whereas ‘***B***’, ‘***C***’, and ‘***D***’ represented the 20 mM, 40 mM, and 80 mM EMS concentrations, respectively. The numbers following each letter corresponded to the sequential numbering of plants within each treatment group. Thus, genotypes from the control (0 mM EMS) were labelled A1–A31, those treated with 20 mM EMS were labelled B1–B48, genotypes exposed to 40 mM EMS were designated C1–C55 and those treated with 80 mM EMS were labelled D1–D56.

The treated seeds were thoroughly washed in running water for 30 minutes to reduce the residual effect of the mutagen sticking to the seed coat. The treated seeds, along with the control, were planted to raise the M_1_ generation, and standard cultural practices were carried out thereafter. The experiment was designed in pots measuring 3.29 m^2^. Spacing used was 0.8m between lines and 0.5 m between rows. One cowpea seed was planted per hole in each pot, at a depth of 3-4 cm.

### Parameters studied in the M_1_ generation

All traits studied were measured and/or counted using standard cowpea descriptor from [28]. The growth, yield-related and yield parameters studied in the M1 generation were grouped as phenological: days to germination (*dtg*), days to first flower (*dff*), days to 50% flowering (*d50f*), days to first harvest (*dfh*), days to 50% mature pod (*d50mp*); and yield-related traits: peduncle length at first harvest (*pel*), number of pods per peduncle (*npp*), number of pods per plant at maturity (*nptm*), number of seeds per pod (*nsp*), number of seeds per plant (*nspt*), pod length (*pdl*), pod width (*pdw*), number of locules (*ln*), percent seed abortion (*psa*). Seed dimensions included seed length (*sdl*), seed width (*sdw*) and seed thickness (*sdt*),while other parameters measured were seed weight (*swg*) and yield per plant (*yld*).

Percentage seed abortion was calculated as;

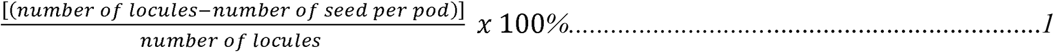

Yield (g) was calculated as:

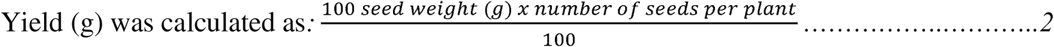

Seed germination was recorded 7 days after planting. Survival plants were recorded at 50% flowering, and germination and survival rate were recorded using the formula by [29]

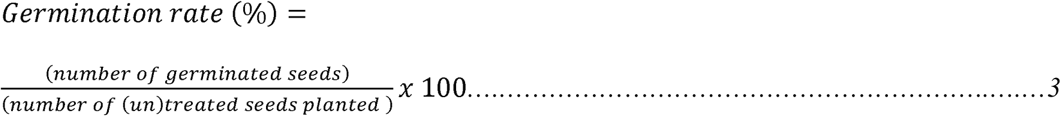

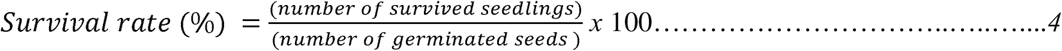

### Statistical analysis

#### Regression seedling germination and survival rate

XLSTAT was also used to generate the linear regression equation of seedling germination and survival rate.

#### Distribution and post hoc contrast analysis for the selection of putative traits and mutants’

Descriptive statistics, one-way ANOVA and post hoc contrast analysis were used to compare treatment effects for each trait in Jamovi 2.7.15.0. Plants within treatments whose values fell outside the control range for each character and were statistically significant from the control were selected as putative mutants.

#### Multivariate analysis for the verification of the selected putative mutant traits and genotypes

Principal Component Analysis (PCA) was performed, accompanied by biplot analysis in Jamovi 2.7.15.0. Principal Component Analysis for quantitative traits was employed to determine the percentage contribution of each trait to total genetic variation. The principal components (PCs) with eigenvalues >1 were selected and traits with coefficients >0.3 were considered to have significantly contributed to the variability. The biplot of computed similarity values was used to visualize relationships among traits across EMS treatment groups and genotypes, to assess the distribution of treatment groups, examine trait vector orientations and identify putative mutants based on their displacement from the control.Cluster analyses were performed using Jamovi 2.7.15.0 to group genotypes within treatments using the Ward D2 hierarchical clustering optimization criterion. A circular dendrogram constructed from computed similarity values was used to visualize relationships among genotypes and traits that differed significantly among treatments, with the resulting clusters interpreted to characterize the phenotypic groupings produced by mutagenic treatment.

## Results

### Percentage germination and survival of M_1_ plants

The results for seed germination and survival rates (%) and linear regression fits for M□ cowpea genotypes subjected to three EMS doses for 6 h are presented in Fig 1. The highest seed germination rate (89.33%) and seedling survival rate (74.67%) were recorded at 80 mM, whereas the lowest germination rate (70.00%) and seedling survival rate (62.00%) were observed in the 0 mM (control) treatment. The percentages of seed germination and seedling survival were in this order: 0 mM < 20 mM < 40 mM < 80 mM. Linear regression analysis revealed a strong positive relationship between EMS concentration and both seed germination and survival rate (Fig 1). For seed germination rate, the regression equation was y = 6.47x + 65.33 with an R^2^ value of 0.94. Seedling survival showed a positive linear relationship with EMS concentration as follows: y = 4.73x + 56.67 with an R^2^ value of 0.90.

**Fig 1:**
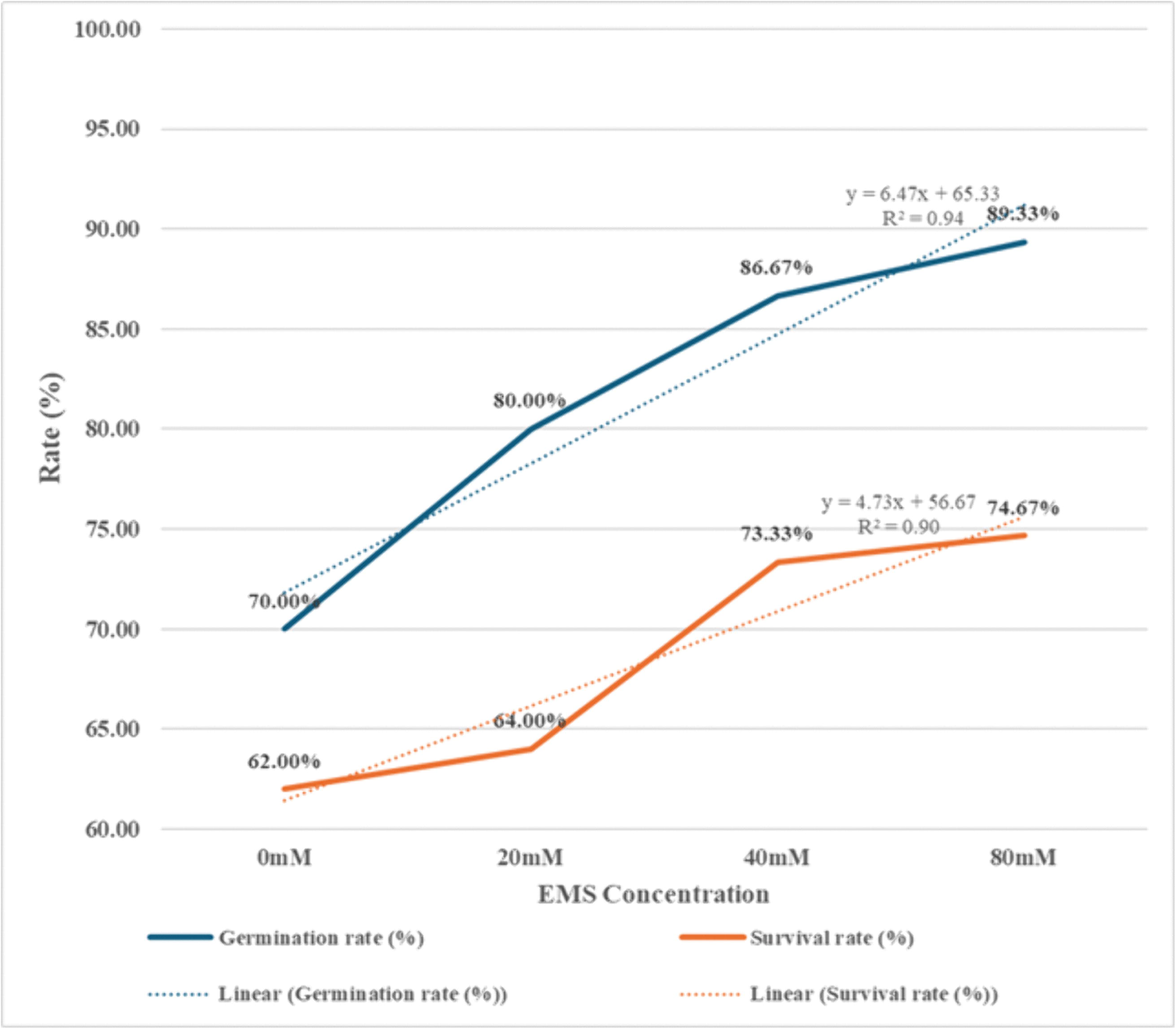
Seed germination and survival rates (%) and linear regression fits for M□ cowpea genotypes subjected to three ethyl methanesulfonate (EMS) doses for 6 h.

### Contrast analysis of traits in the M_1_ generation

#### Phenological traits

***Days to germination* (*dtg*)** differed significantly among treatments (P < 0.001). The control (0 mM) germinated earliest, with a mean of 3.45 ± 0.13 days and a range of 2–5 days. All EMS treatments delayed germination, with the most substantial effect at 20 mM, while 40 and 80 mM showed delayed but comparable responses. The frequency of putative mutants was 50.0% at 20 mM, 29.10% at 40 mM, and 30.36% at 80 mM (Table 1). Selected putative mutants can be found in S1 Table.

**Table 1:** Mean, variability, mutant frequency, and pairwise significance of phenological traits in the M□ generation of cowpea under different EMS concentrations.

| Trait | N | Trt | Mean±se | %CV | Min | Max | Freq | P | T-values of significant pairwise comparisons |  |  |  |
| --- | --- | --- | --- | --- | --- | --- | --- | --- | --- | --- | --- | --- |
|  |  |  |  |  |  |  |  |  | 0mM | 20mM | 40mM | 80mM |
| dtg | 31 | 0mM | 3.45±0.13<sub>a</sub> | 20.90 | 2 | 5 |  | *** |  | -6.38*** | -4.53*** | -4.50*** |
|  | 48 | 20mM | 5.38±0.20<sub>b</sub> | 25.30 | 3 | 7 | 50.00 |  | -6.38*** |  |  |  |
|  | 55 | 40mM | 4.78±0.19<sub>b</sub> | 29.20 | 3 | 7 | 29.10 |  | -4.53*** |  |  |  |
|  | 56 | 80mM | 4.77±0.19<sub>b</sub> | 29.70 | 3 | 7 | 30.36 |  | -4.50*** |  |  |  |
| dff | 31 | 0mM | 37.87±0.77<sub>a</sub> | 11.30 | 30 | 45 |  | *** |  | -5.00*** | -5.40*** | -4.97*** |
|  | 48 | 20mM | 44.69±0.91<sub>b</sub> | 14.10 | 33 | 54 | 51.17 |  | -5.00*** |  |  |  |
|  | 55 | 40mM | 45.04±0.86<sub>b</sub> | 14.10 | 34 | 55 | 47.27 |  | -5.40*** |  |  |  |
|  | 56 | 80mM | 44.46±0.79<sub>b</sub> | 13.30 | 33 | 55 | 41.07 |  | -4.97*** |  |  |  |
| d50f | 31 | 0mM | 57.84±0.92<sub>a</sub> | 8.90 | 48 | 68 |  | *** |  | -4.51*** | -4.98*** | -4.72*** |
|  | 48 | 20mM | 64.67±0.99<sub>b</sub> | 10.70 | 50 | 75 | 37.50 |  | -4.51*** |  |  |  |
|  | 55 | 40mM | 65.18±0.97<sub>b</sub> | 11.00 | 49 | 80 | 40.00 |  | -4.98*** |  |  |  |
|  | 56 | 80mM | 64.79±0.85<sub>b</sub> | 9.80 | 53 | 80 | 28.57 |  | -4.72*** |  |  |  |
| dfh | 31 | 0mM | 49.35±1.15<sub>a</sub> | 13.00 | 40 | 67 |  | *** |  | -6.93*** | -6.93*** | -6.04*** |
|  | 48 | 20mM | 60.17±0.94<sub>b</sub> | 10.80 | 48 | 72 | 18.75 |  | -6.93*** |  |  |  |
|  | 55 | 40mM | 58.93±0.97<sub>b</sub> | 12.10 | 38 | 72 | 9.100 |  | -6.93*** |  |  |  |
|  | 56 | 80mM | 58.50±0.91<sub>b</sub> | 11.70 | 43 | 72 | 12.50 |  | -6.04*** |  |  |  |
| d50mp | 31 | 0mM | 79.48±1.41<sub>a</sub> | 9.90 | 66 | 94 |  | *** |  | -5.40*** | -4.75*** | -4.59*** |
|  | 48 | 20mM | 90.17±1.37<sub>b</sub> | 10.50 | 68 | 107 | 39.58 |  | -5.40*** |  |  |  |
|  | 55 | 40mM | 88.65±1.09<sub>b</sub> | 9.10 | 68 | 106 | 25.45 |  | -4.75*** |  |  |  |
|  | 56 | 80mM | 88.32±1.16<sub>b</sub> | 9.80 | 69 | 110 | 23.21 |  | -4.59*** |  |  |  |

Note. Means sharing the same superscript letter within a trait are not significantly different according to Tukey's HSD test ( $P \leq 0.05$ ). Trt= Treatment, N=Sample size, Mean ± SE = Mean plus or minus standard error, %CV= Percentage coefficient of variation, Min= minimum, Max= maximum, Freq= Percentage frequency of putative mutants; P=Probability value. Traits: dtg = days to germination, dff = days to first flower, d50f = days to 50% flowering, dfh = days to first harvest, d50mp = days to 50% mature pods, \*=significance at 5% level, \*\*=significance at 1% level, \*\*\*=significance at 0.1% level

***Days to first flower (dff)*** were significantly affected by EMS across treatments (P < 0.001). *DFF* was the earliest in the control, with a mean of 37.87 ± 0.77 days and a range of 30 to 45 days, and was progressively delayed with increasing EMS concentration, reaching a maximum mean of 45.04 ± 0.86 days at 40 mM. Mutant frequencies declined from 51.17% at 20 mM to 41.07% at 80 mM (Table 1). Selected putative mutants can be found in S1 Table.

***Days to 50% flowering (d50f)*** were significantly affected by EMS across treatments (P < 0.001). *D50f* increased from a control mean of 57.84 ± 0.92 days, with values ranging from 48 to 68 days, to mean values of approximately 65 days across EMS treatments, with broader range of 49–80 days. Corresponding mutant frequencies were 37.50%, 40.00%, and 28.57% at 20, 40, and 80 mM, respectively (Table 1). Selected putative mutants can be found in S1 Table.

***Days to first harvest (dfh)*** were the earliest in the control, with a mean of 49.35 ± 1.15 days and a range of 40–67 days and were significantly (P < 0.001) delayed in all EMS treatments. Mean dfh increased to 60.17 ± 0.94 days at 20 mM, 58.93 ± 0.97 days at 40 mM, and 58.50 ± 0.91 days at 80 mM. Mutant frequencies were 18.75% at 20 mM, 9.10% at 40 mM, and 12.50% at 80 mM (Table 1). Selected putative mutants can be found in S1 Table.

A similar trend was observed for days to ***50% maturity of pods (d50mp)***, with the control recording the earliest maturity at 79.48 ± 1.41 days, and values ranging from 66 to 94 days. EMS treatments significantly (P<0.001) delayed pod maturity compared with the control, with mean d50mp values of 90.17 ± 1.37 days at 20 mM, 88.65 ± 1.09 days at 40 mM, and 88.32 ± 1.16 days at 80 mM. The frequency of selected mutants for d50mp was 39.58%, 25.45%, and 23.21% at 20, 40, and 80 mM, respectively (Table 1). Selected putative mutants can be found in S*1* Table.

#### Yield-related and yield traits

***Pod length (pdl)*** was highest in the control (0 mM), with a mean of 14.65 ± 0.31 cm and a range of 11.55–17.90 cm, whereas 80mM recorded the lowest mean pdl of 13.52±0.28 cm, which was significantly different (P=0.008) from the control. Based on deviations from the control range, the frequency of selected mutants for pdl was 10.42%, 9.10%, and 21.43% in 20 mM, 40 mM, and 80 mM, respectively (Table 2). Selected putative mutants can be found in S1 Table.

**Table 2:**
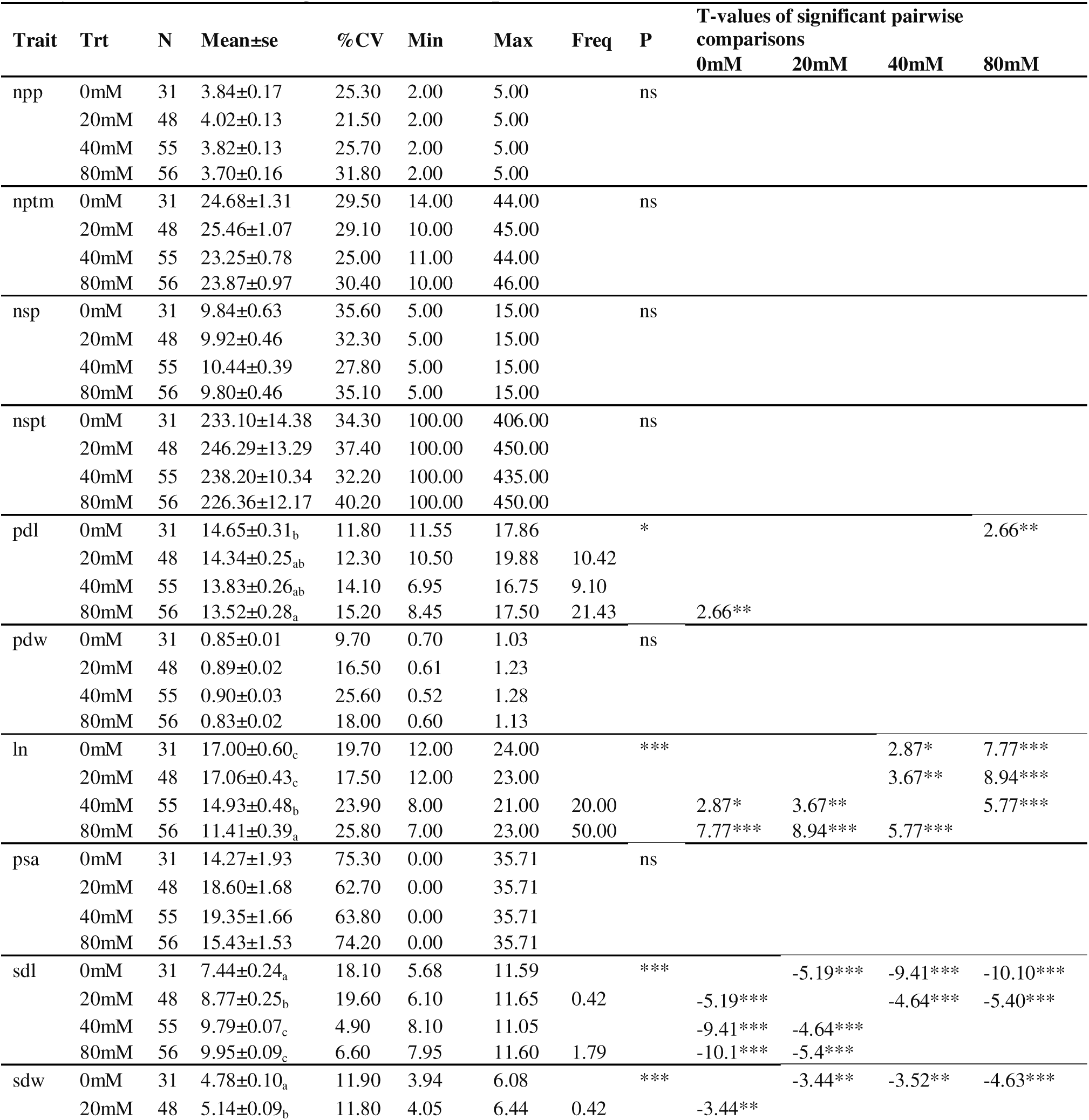
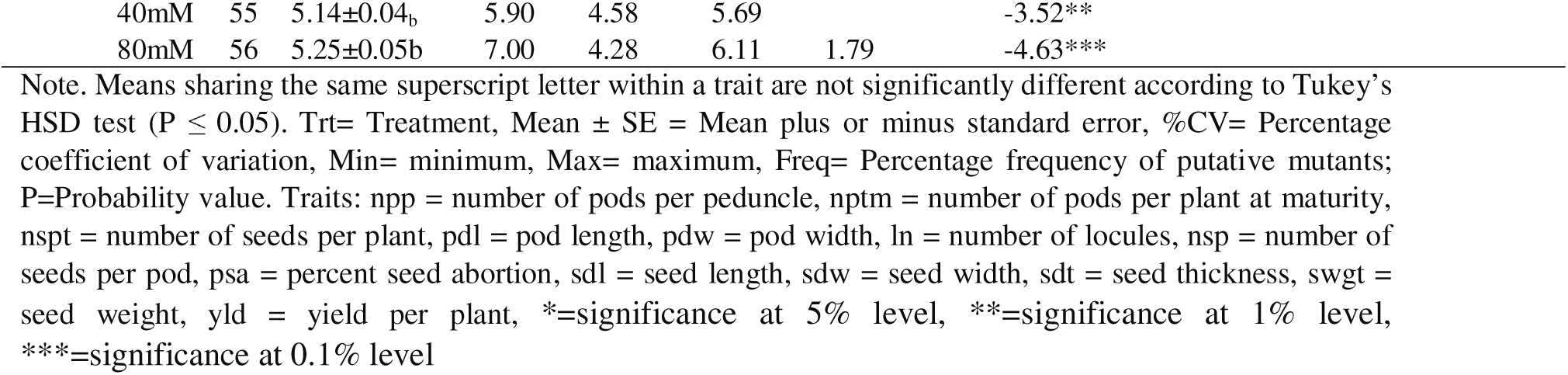
Mean, variability, mutant frequency, and pairwise significance of yield-related and yield traits in the M□ generation of cowpea under different EMS concentrations.

**Table 2:** Mean, variability, mutant frequency, and pairwise significance of yield-related and yield traits in the M□ generation of cowpea under different EMS concentrations.

| Trait | Trt | N | Mean±se | %CV | Min | Max | Freq | P | T-values of significant pairwise comparisons | | | |
| --- | --- | --- | --- | --- | --- | --- | --- | --- | --- | --- | --- | --- |
|  |  |  |  |  |  |  |  |  | 0mM | 20mM | 40mM | 80mM |
| sdt | 0mM | 31 | 4.09±0.07<sub>a</sub> | 9.80 | 3.29 | 4.92 |  | *** |  | -2.82* |  |  |
|  | 20mM | 48 | 4.35±0.08<sub>b</sub> | 12.20 | 3.31 | 5.19 | 12.50 |  | -2.82* |  | 4.92*** | 3.35** |
|  | 40mM | 55 | 3.95±0.04<sub>a</sub> | 8.50 | 3.43 | 5.59 | 1.82 |  |  | 4.92*** |  |  |
|  | 80mM | 56 | 4.08±0.05<sub>a</sub> | 8.90 | 3.41 | 4.94 | 1.79 |  |  | 3.35** |  |  |
| swg | 0mM | 31 | 21.03±0.34<sub>a</sub> | 9.10 | 17.82 | 24.87 |  | *** |  | -6.91*** | -6.18*** | -2.92* |
|  | 20mM | 48 | 24.86±0.34<sub>c</sub> | 9.40 | 17.81 | 29.99 | 54.17 |  | -6.91*** |  |  | 4.77*** |
|  | 40mM | 55 | 24.37±0.35<sub>c</sub> | 10.70 | 17.70 | 28.39 | 49.10 |  | -6.18*** |  |  | 3.87*** |
|  | 80mM | 56 | 22.60±0.33<sub>b</sub> | 11.00 | 17.35 | 28.68 | 16.07 |  | -2.92* | 4.77*** | 3.87*** |  |
| yld | 0mM | 31 | 48.79±3.01<sub>a</sub> | 34.40 | 20.61 | 85.68 |  | * |  | -2.61* |  |  |
|  | 20mM | 48 | 61.19±3.34<sub>b</sub> | 37.80 | 18.70 | 125.44 | 16.67 |  | -2.61* |  |  |  |
|  | 40mM | 55 | 58.02±2.59<sub>ab</sub> | 33.10 | 19.97 | 105.43 | 12.72 |  |  |  |  |  |
|  | 80mM | 56 | 51.16±2.87<sub>ab</sub> | 42.00 | 18.45 | 111.85 | 10.71 |  |  |  |  |  |

Note. Means sharing the same superscript letter within a trait are not significantly different according to Tukey's HSD test ( $P \leq 0.05$ ). Trt= Treatment, Mean ± SE = Mean plus or minus standard error, %CV= Percentage coefficient of variation, Min= minimum, Max= maximum, Freq= Percentage frequency of putative mutants; P=Probability value. Traits: npp = number of pods per peduncle, nptm = number of pods per plant at maturity, nspt = number of seeds per plant, pdl = pod length, pdw = pod width, ln = number of locules, nsp = number of seeds per pod, psa = percent seed abortion, sdl = seed length, sdw = seed width, sdt = seed thickness, swgt = seed weight, yld = yield per plant, \*=significance at 5% level, \*\*=significance at 1% level, \*\*\*=significance at 0.1% level

***Number of locules per pod (ln)*** differed significantly among EMS treatments (P < 0.001). The control recorded a mean of 17.00 ± 0.60, with a range of 12 to 24 locules. Higher EMS concentrations resulted in a significant reduction in ln: 14.93 ± 0.48 locules at 40 mM and 11.41 ± 0.39 locules at 80 mM. Pairwise comparisons showed that only the 80 mM treatment differed significantly (P < 0.001). From both the control and all other treatments, Phenotypic variability increased with EMS dose, as reflected by a rise in the coefficient of variation from 17.5% at 20 mM to 25.8% at 80 mM. Based on deviations from the control range, the frequency of putative mutants for ln reached 20.0% at 40 mM and 50.0% at 80 mM (Table 2). Selected putative mutants can be found in S1 Table.

***Seed length*** *(**sdl**)* varied significantly across EMS treatments (P < 0.001). The control recorded the lowest mean seed length, with a value of 7.44 ± 0.24 mm, with a range of 5.68 to 11.59 mm and a CV of 18.1%. EMS treatment groups resulted in a progressive increase in seed length, with means of 8.77 ± 0.25 mm at 20 mM, 9.79 ± 0.07 mm at 40 mM, and 9.95 ± 0.09 mm at 80 mM. Pairwise comparisons showed that all EMS treatments differed significantly from the control (P < 0.001), while the 40 mM and 80 mM treatments did not differ from each other. Based on deviations from the control range, mutant frequencies were low for seed length recorded at 0.42% for 20 mM and 1.79% for 80 mM (Table 2). Selected putative mutants can be found in S1 Table.

***Seed width (sdw)*** was also significantly (P < 0.001) affected by EMS treatment. The control population had a mean width of 4.78 ± 0.10 mm, ranging from 3.94 to 6.08 mm, with a CV of 11.9%. All EMS treatments differed significantly from the control (P < 0.001). Variability declined at higher EMS doses, with CV values as low as 5.9% at 40 mM. Frequencies of putative mutants increased with EMS concentration, reaching 0.42% at 20 mM and 1.79% at 80 mM (Table 2). Selected putative mutants can be found in S1 Table.

***Seed thickness (sdt)*** differed significantly among treatments (P < 0.001). The control recorded a mean thickness of 4.09 ± 0.07 mm with values ranging from 3.29 to 4.92 mm. Thickness increased significantly (P=0.027) at 20 mM with a mean of 4.36 ± 0.08 mm. Pairwise comparisons indicated significant differences between 20 mM and both 40 mM (P < 0.001) and 80 mM (P = 0.005). Mutant frequencies for seed thickness were highest at 20 mM (12.5%) but remained low at 40 mM (1.82%) and 80 mM (1.79%) (Table 2). Selected putative mutants can be found in S1 Table.

***100-Seed weight (swg)*** showed highly significant variation among EMS treatments (P < 0.001). The control recorded a mean of 21.03 ± 0.34 g, with a range of 17.82 to 24.87 g. EMS treatment increased swg, with the highest means observed at 20 mM (24.86 ± 0.34 g) and 40 mM (24.37 ± 0.35 g), both of which differed significantly from the control (P < 0.001). The 80 mM treatment weighed 22.60 ± 0.33 g and showed an intermediate response, differing considerably from both the control (P = 0.02) and lower EMS doses (P < 0.001). Frequencies of putative mutants were highest at 20 mM (54.17%), followed by 40 mM (49.1%) and 80 mM (16.07%) (Table 2). Selected putative mutants can be found in S1 Table.

***Yield per plant (yld)*** differed significantly among treatments (P = 0.018). The control population recorded the lowest mean yield of 48.79 ± 3.01 g, with a range of 20.61 to 85.68 g. Yield increased significantly (P = 0.047) at 20 mM, with a mean of 61.19 ± 3.34 g, which differed from the control (P = 0.047). The 40 mM (58.02 ± 2.59 g) and 80 mM (51.16 ± 2.87 g) treatments, which exhibited intermediate yields, did not differ significantly from either the control or the 20 mM treatment. Phenotypic variability was high across treatments, with a CV of 33.10–42%, and mutant frequencies declined with increasing EMS concentration, from 16.67% at 20 mM to 10.71% at 80 mM (Table 2). Selected putative mutants can be found in S1 Table.

### Multivariate Analysis of Phenological and Yield-Related Traits

#### PCA Biplot of phenological, yield-related and yield traits

##### Variance Explained by Principal Components

Principal component analysis (PCA) of the traits and genotypes of plant observations across EMS treatments yielded 8 principal components with eigenvalues < 1 (S2 Table). The first four principal components accounted for a total variability greater than 50.60% in all treatments. The first two principal components (Dim1 and Dim2) together explained 32.8% of the total phenotypic variation, with PC1 accounting for 17.7% and PC2 accounting for 15.1% (Fig 2, S2 Table) with an eigenvalues of 3.18 and 2.71 respectively.

**Fig 2:**
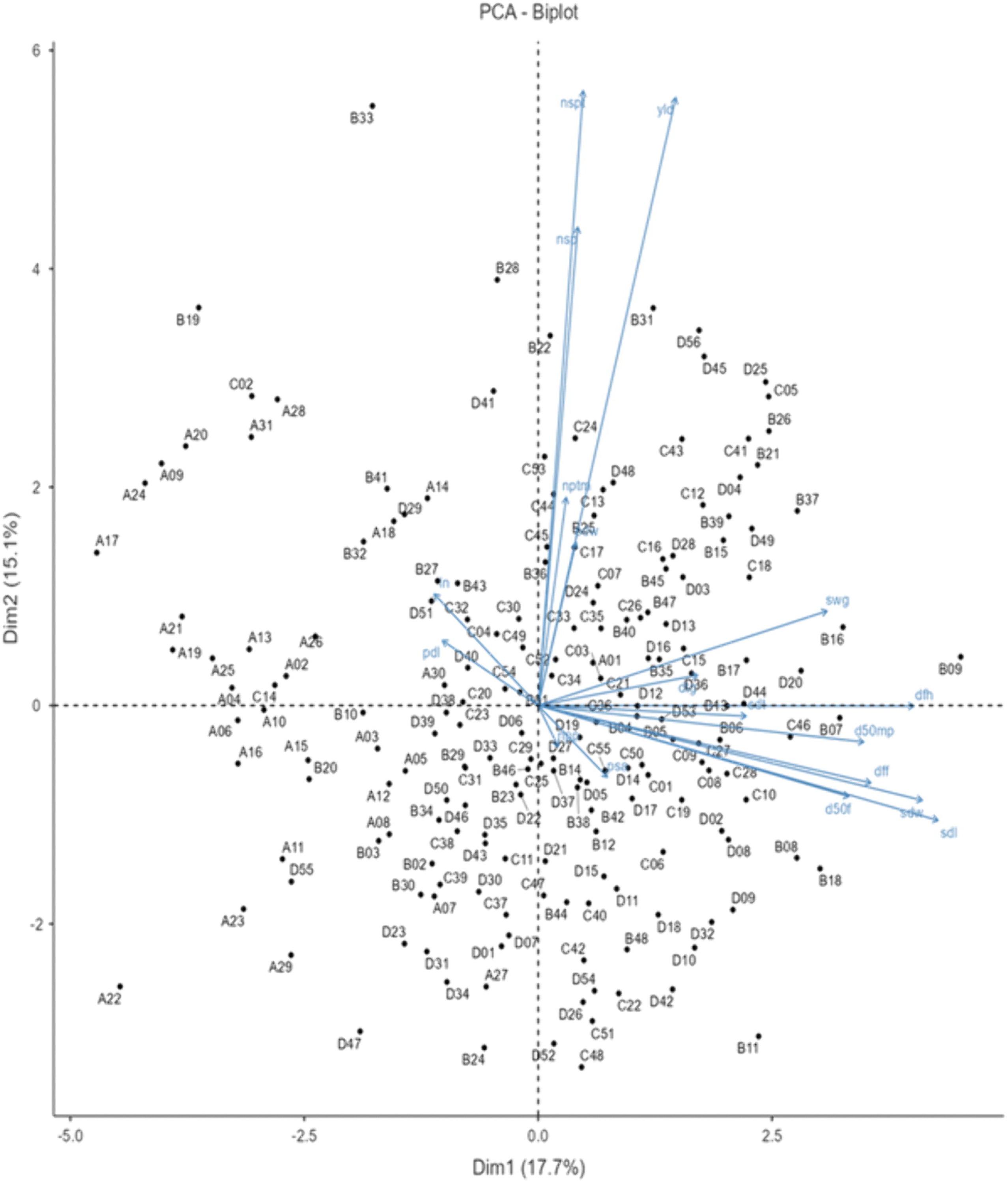
PCA biplot of cowpea genotypes and traits based on PC1 and PC2. (a)Genotypes treated with 0-, 20-, 40- and 80-mM EMS are labelled A01–A31, B01–B48, C01–C55 and D01–D56, respectively. (b) Vectors indicate trait loadings, and points represent genotype distribution. (c) dtg= days to germination, dff = days to first flower, d50f = days to 50% flowering, dfh = days to first harvest, d50mp = days to 50% mature pods, npp = number of pods per peduncle, nptm = number of pods per plant at maturity, nspt = number of seeds per plant, pdl = pod length, pdw = pod width, ln = number of locules, nsp = number of seeds per pod, psa = percent seed abortion, sdl = seed length, sdw = seed width, sdt = seed thickness, swgt = seed weight, yld = yield per plant

##### Distribution of EMS Treatment Groups in the PCA Biplot

The PCA biplot (Fig 2) plotted genotype scores on PC1 (Dim1, 17.7%) against PC2 (Dim2, 15.1%). Control genotypes (A01–A31, 0 mM) clustered predominantly on the negative side of PC1, spanning approximately −5.0 to −0.5, with limited spread along PC2. Genotypes from the 20 mM treatment (B01–B48) occupied a broader area, with partial overlap with the control cluster and moderate displacement toward positive PC1 values. Genotypes from the 40 mM treatment (C01–C55) were more widely dispersed across both axes, with a concentration in the central to positive PC1 region. Genotypes from the 80 mM treatment (D01–D56) showed the widest distribution across the biplot, with the greatest displacement toward positive PC1 and the greatest scatter along PC2 of any treatment group.

##### Biplot Trait Vector Orientations

Vectors for *dff*, *d50f*, *dfh*, and *d50mp* projected horizontally toward the positive PC1 axis. Vectors for sdl and sdw were oriented toward the lower-right quadrant (positive PC1, negative PC2). The *swg* vector projected toward the positive region of both PC1 and PC2. Vectors for *yld*, *nspt*, and *nsp* were directed upward toward positive PC2, with *yld* and nspt nearly collinear. The *nptm* vector was oriented downward toward negative PC2. Vectors for *pdl* and *ln* projected toward the upper-central region of the biplot. The angle between the *sdl*/*sdw* vectors and the *yld*/*nspt*/*nsp* vectors was approximately perpendicular (Fig 2).

##### Identification and Selection of Putative Mutants from the Biplot

Based on positional alignment with trait vectors of agronomic interest, three classes of putative mutants were identified in Fig 2. Eleven genotypes positioned in the upper biplot region with positive PC2 scores, in alignment with the *yld*, *nsp*, *nspt*, and *swg* vectors, were classified as high-yielding putative mutants: **B33, B28, B22, B31, D56, D45, D25, C5, C41, B26, and B21**. Eleven genotypes positioned along the negative PC2 axis in alignment with the *sdl* and *sdw* vectors, showing above-average seed dimensions with reduced seed number and yield, were classified as large-seed putative mutants: **D23, D7, D34, D47, C37, C39, C40, C42, B18, B11** and **B8**.

Two genotypes positioned on the extreme negative PC1 side, in the direction opposite to the phenological delay vectors, were classified as early-phenology putative mutants: **B19** and **C2**. Four genotypes **B9, B7, C46 and D20** were positioned at the extreme positive PC1 end with weak alignment to yield vectors and were classified as severely delayed and low performing. These genotypes were not selected for advancement but were retained as extreme phenotypic reference individuals.

#### Cluster analysis of phenological and yield-related traits

Hierarchical cluster analysis of 18 phenological and yield-related traits across 190 M□ cowpea genotypes using Ward’s linkage method resolved six distinct clusters (Fig 3, S1 Table). The six clusters contained 64, 24, 31, 20, 40, and 9 genotypes, respectively.

**Fig 3:**
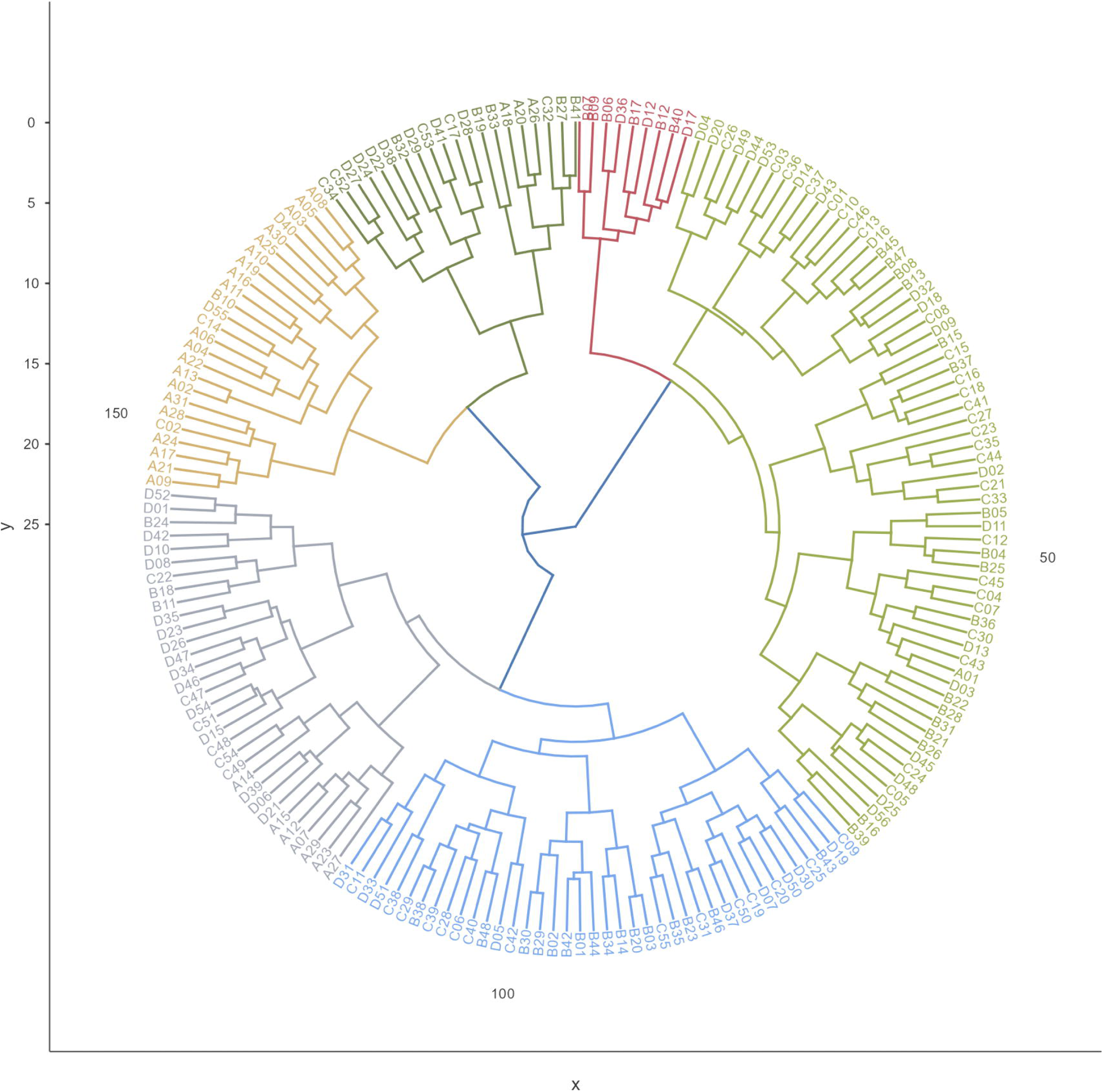
Dendrogram of hierarchical cluster analysis of MD cowpea genotypes based on quantitative traits using Ward’s linkage method. (a)Genotypes treated with 0-, 20-, 40-, and 80-mM EMS are labelled A, B, C, and D, respectively. (b) Six clusters are delineated by colour: Cluster 1 (green, n = 64); Cluster 2 (brown, n = 24); Cluster 3 (grey, n = 31); Cluster 4 (forest green, n = 20); Cluster 5 (blue, n = 40); Cluster 6 (red, n = 9).

##### Cluster 1 (n = 64)

Cluster 1 contained predominantly EMS-treated genotypes, with 35.42%, 49.10%, and 33.93% of the 20 mM, 40 mM, and 80 mM treatment groups represented (Fig 3, S1 Table). Only one control genotype (A01) was assigned to this cluster, accounting for 3.22% of the 0 mM population. Days to 50% flowering ranged from 53 to 80 days across cluster members, with the longest delays recorded in C41 and C43. Days to 50% pod maturity ranged from 69 to 109 days, with the most extended values recorded in D43 and C08. Seed weight ranged from 18.82 g (D04) to 28.68 g (D56). Yield per plant ranged from 31.45 g (C37) to 111.85 g (D56). Putative mutants selected from this cluster included D56 (111.85 g), D45 (*106.09 g*), D25 (*102.66 g*), B31 (*107.22 g*), B28 (*103.50 g*), B22 (*88.96 g*), C41 (*90.96 g*) and C5 (*105.43 g*).

##### Cluster 2 (n = 24)

Cluster 2 was dominated by control genotypes, which accounted for 61.29% of the 0 mM population (Fig 3, S1 Table). Representation from EMS treatments was low: 2.08% (20 mM), 3.64% (40 mM), and 3.57% (80 mM). Genotypes in this cluster had the earliest germination, flowering, and maturity values across the full population, with yield levels comparable to the control mean and low within-cluster variability. EMS-treated genotypes assigned to this cluster included C02, with days to first harvest of 38 days, and C14, with days to first harvest of 39 days. Both were identified as early-phenology putative mutants.

##### Cluster 3 (n = 31)

Cluster 3 contained genotypes primarily from the high-dose EMS treatments, with 28.57% of the 80 mM population represented (Fig 3, S1 Table). Genotypes in this cluster showed delayed phenological development, reduced locule number, and low yield per plant. Representative genotypes included B7, B9, C46, and D20, all of which showed negative deviation from the control range across multiple yield traits. These genotypes were not selected for advancement.

##### Cluster 4 (n = 20)

Cluster 4 included genotypes from all three EMS treatment groups: 10.42% of 20 mM, 9.09% of 40 mM, and 12.50% of 80 mM (Fig 3, S1 Table). Genotypes in this cluster combined relatively early days to first flower and days to 50% flowering with extended days to first harvest and days to 50% mature pods. Seed weight and yield per plant were elevated relative to the control in several members. The highest-yielding genotype in this cluster was B33 (125.44 g), followed by B19 (93.59 g). Both substantially exceeded the control population range for yield per plant while retaining comparatively early flowering phenology — B19 at 36 days to first flower and B33 at 37 days.

##### Cluster 5 (n = 40)

Cluster 5 comprised exclusively EMS-treated genotypes, with 33.33% of the 20 mM population, 22.27% of the 40 mM population, and 16.07% of the 80 mM population assigned to this cluster (Fig 3, S1 Table). No control genotypes were present. Genotypes in this cluster showed moderately delayed phenological development and high seed weight but reduced locule number per pod, which was associated with lower yield per plant relative to the control mean.

##### Cluster 6 (n = 9)

Cluster 6 was the smallest cluster, containing 12.50% of the 20 mM population and 5.36% of the 80 mM population (Fig 3, S1Table). No control or 40 mM genotypes were present. Genotypes in this cluster showed moderately delayed phenological development, large seed dimensions, and high seed weight. Seed number per pod was relatively low, but total yield per plant was elevated, driven by seed weight. Putative mutants selected from this cluster included B07 (yld = 67.18 g, swg = 29.99 g) and D36 (yld = 74.69 g, swg = 28.29 g), both of which exceeded the control mean for seed weight.

## Discussion

### Percentage germination and survival of M1 plants

The EMS concentrations of 20, 40, and 80 mM and the fixed 6 h exposure duration were selected to provide a graded mutagenic treatment range while maintaining sufficient plant survival for evaluation, consistent with previous studies showing that EMS response depends on both concentration and exposure time [13,30,31]. In the present study, LD□□ was not achieved, as seed germination and seedling survival increased with increasing EMS concentration. Although most EMS mutagenesis studies report a decline in germination and survival with increasing dose [16,32–34], several studies have documented non-linear or hormetic responses, where low to moderate EMS doses stimulate germination and early growth before inhibitory effects appear at higher concentrations. In legumes, responses similar to the present study have been reported. For instance, in cowpea relatives, [35] observed in faba bean (*Vicia faba* L.) that six varieties exhibited higher germination and survival at 0.05% EMS compared to the control, followed by a decline at higher concentrations. This hormesis-like response mirrors the increasing germination trend observed in the present study. Similarly, in soybean (*Glycine max* L.), [36] reported that EMS treatments at 0.1–0.2% improved germination and early seedling growth relative to untreated seeds, with inhibitory effects only becoming evident at higher doses. In cucumber (*Cucumis sativus* L.), [37] demonstrated that post-treatment handling influenced germination outcomes, with a sodium thiosulfate rinse yielding higher germination (84.4%) than a water rinse (80.0%) at 0.5% EMS, indicating that detoxification and recovery processes can enhance germination performance. Likewise, in tef (*Eragrostis tef*), [38] reported that a treatment combination of 2.50% EMS for 4 h resulted in significantly higher germination (77.1%) compared to lower doses, emphasizing the importance of genotype × dose × exposure-time interactions. In barley (*Hordeum vulgare* L.), [39] also noted a non-monotonic response, where low EMS exposure improved germination relative to some treatments, although optimal survival occurred at intermediate doses. Collectively, these studies support the interpretation that the increasing germination and survival rates observed with rising EMS concentration in the present study may reflect a hormetic or priming-like effect, rather than a deviation from expected mutagenic behavior. This reinforces the role of genotype-specific responses and treatment conditions in shaping EMS-induced physiological outcomes.

Our results differs from the findings of [16, 32–34], [32] reported an apparent dose-dependent reduction in germination and survival in the M□ generation of cowpea varieties *Pusakomal* and *V-240 (Rambha)* following EMS treatments at 0.1–0.3%, with control germination exceeding 95% and higher doses approaching the LD□□ threshold. Similarly, [14] observed a strong negative linear relationship between EMS concentration and germination in cowpea and identified an optimal LD□□ dose at 0.4% EMS. The contrast observed in this current study was that germination and survival were lowest in the control. It increased with EMS concentration, indicating that LD□□ expression is highly dependent on genotype, EMS dose range, and exposure duration. Unlike previous studies that employed percentage-based EMS concentrations and different treatment protocols, the present study used millimolar EMS concentrations with a fixed 6-h exposure, which may have altered the physiological response. In addition, the relatively low germination and survival recorded in the control suggest baseline establishment constraints that may have obscured EMS-induced lethality. These findings indicate that the absence of an LD□□ endpoint in the present study does not imply insufficient mutagenic effect. Instead, it underscores the importance of genotype × dose × exposure-time interactions in EMS mutagenesis. [40,41]. Such responses have been linked to priming-like effects, where mild stress exposure improves germination performance [40].

### Contrast analysis of traits in the M_1_ generation

#### Effects of EMS Mutagenesis on Phenological traits

##### Effect of EMS on Days to Germination

The present study revealed that EMS treatment significantly delayed days to germination (*dtg*) in the M□ generation of cowpea (*Vigna unguiculata* (L.) Walp). The control seeds germinated earliest, while all EMS-treated groups exhibited delayed germination. This pattern of EMS-induced germination delay is consistent with the well-established understanding that chemical mutagens impose physiological stress during seed imbibition, activating DNA repair mechanisms that temporarily arrest the cell cycle until lesions are repaired [31]. The alkylating activity of EMS, which produces O□-ethylguanine and other base modifications, necessitates repair via base excision repair pathways before the resumption of cell division in the embryonic axis [21]. This repair process delays radicle emergence, resulting in a measurable increase in days to germination.

Comparable delays in days to germination have been documented in other crops. Similarly, research on marigold (*Tagetes* spp.) demonstrated that higher EMS concentrations and longer exposure times reduced germination rate and delayed progress, although the effect was largely inhibitory rather than showing a peak delay at a lower dose [42]. The high frequency of putative mutants for *dtg* at 20 mM (50.0%) compared to 40 mM (29.1%) and 80 mM (30.4%) aligns with the observation that lower EMS doses often yield a higher proportion of viable mutants because they cause less overall lethality and permit the expression of a wider range of phenotypic variation [23,24]. In cowpea, [13] similarly found that lower EMS doses produced useful variability while higher doses reduced survival and limited the number of recoverable mutants. The reduction in mutant frequency at higher concentrations in our study may reflect the increased lethality and physiological damage that suppress the recovery of germination-related mutants.

##### Effects of EMS on Days to First Flower

In the present study, EMS treatment at all three concentrations caused a highly significant delay in *dff* relative to the control. Pairwise comparisons confirmed that all EMS treatments differed significantly from the control, while no significant differences were detected among the three EMS concentrations themselves. This pattern suggests a threshold effect, where even the lowest dose was sufficient to disrupt normal floral transition, with no further meaningful delay introduced at higher concentrations. The delayed onset of flowering observed here aligns with findings reported in other EMS mutagenesis studies on cowpea. [26] documented early and late maturity as among the viable mutant classes induced by EMS in *Vigna unguiculata*, noting that EMS can modify important components of plant cells and affect the morphology, anatomy, biochemistry, and physiology of plants differentially depending on concentration level. Similarly, [14] reported effects of EMS, DES, and SA on quantitative traits of cowpea in the M□ generation, providing early evidence that flowering-related traits are susceptible to chemical mutagenesis in this species.

The mechanism underlying EMS-induced delays in flowering time is most likely attributable to point mutations disrupting key genes involved in floral transition pathways. [31] noted that M□ plants subjected to EMS mutagenesis have shown variation in flower organs and delayed flowering, consistent with the disruption of genes regulating the floral transition, and that such phenotypic changes in flower-related traits may reflect the broad mutagenic spectrum of EMS across multiple developmental pathways. In cowpea specifically, flowering time is known to be a quantitatively inherited and environmentally sensitive trait. [43] demonstrated that time to flowering in cowpea is strongly influenced by photoperiod, with photoperiod-sensitive genotypes under long natural days exhibiting considerable delays in floral development. EMS-induced mutations affecting photoperiod perception genes or floral integrators such as *GIGANTEA*, *CONSTANS*, or *FLOWERING LOCUS T* homologs could therefore plausibly extend the time to first flower, even at relatively low mutagen concentrations.

The mutation frequencies observed for days to first flower as 51.17%, 47.27%, and 41.07% for 20, 40, and 80 mM, respectively showed a declining trend with increasing EMS concentration. This is consistent with the well-documented inverse relationship between mutagen dose and the proportion of viable, phenotypically expressed mutations, where higher doses tend to induce increasingly lethal or phenotypically cryptic mutations rather than productive, viable phenotypic variants. [44] reported a similar pattern in cowpea var. Arka Garima, where a linear decline in morphological mutation frequency was observed with increasing EMS concentration, suggesting that higher doses induce proportionally more deleterious rather than viable mutations.

##### Effects of EMS on days to 50% flowering

In this study, we observed that the pairwise comparisons confirmed that all mutagenized populations differed significantly from the control but not from each other, reinforcing the threshold dose interpretation. The coefficient of variation was relatively low across all groups (8.9–11%), indicating that although the mean was shifted by EMS treatment, population-level variation in the timing of 50% flowering remained moderate. Notably, the range of days to 50% flowering was broader in the EMS treatments (50–80 days) than in the control (48–68 days), reflecting the expanded phenotypic diversity generated by mutagenesis. These findings are in agreement with results reported from induced mutagenesis experiments in related *Vigna* species and cowpea populations. For instance, [45] evaluated M□ Tswana cowpea mutant lines across two seasons reported significant variation in days to 50% flowering, with the Tswana control consistently flowering earlier than many of the mutant lines and noted that delayed flowering in mutagenized populations may be attributed to genetic changes created by the mutation that affect normal flowering mechanisms.

In EMS-mutagenized cowpea (cv. *Asontem*) evaluated in M□ populations, [46] similarly reported wide distribution of days to flowering among mutagenized individuals compared to the wild type, demonstrating that EMS is effective in generating significant phenotypic variation in flowering-related quantitative traits in cowpea. The parallel shifts in both days to first flower and days to 50% flowering across all EMS treatments suggested that EMS disrupted the floral initiation process rather than the speed of floral development once initiated. If the mutagen had primarily affected floral development after initiation, one would expect a disproportionate shift in days to 50% flowering without a corresponding shift in days to first flower. The approximately equal delays of 6–7 days in both traits across all treatments instead point to a disruption of the upstream signaling or perception pathways governing the transition from vegetative to reproductive growth. Research on contrasting cowpea genotypes by [47] established that delays in flowering are primarily driven by delays in the first appearance of floral buds rather than in bud development itself, supporting the interpretation that the mutagenic effects observed here act at or upstream of bud initiation.

The observations in this study were made in the M□ generation, where induced mutations were expected to be heterozygous, and chimerism may confound phenotypic expression. [46] noted that during the M□ generation, only dominant or co-dominant mutations are easily detected by phenotypic observation, as the functional mutations induced by EMS through point mutation have a greater chance of being dominant or co-dominant.

The delayed flowering documented in this study likely reflected the expression of dominant or semi-dominant mutation effects on floral pathway genes, while recessive mutations which may represent early or altered flowering phenotypes of greater breeding value will only become apparent upon selfing and selection in the M□ and subsequent generations.

The delay in flowering under stronger mutagenesis is a common phenomenon, often attributed to physiological stress or to disruption of genes controlling flowering time [48–51].[49] did not directly measure flowering time, but their M□ mutants showed overall physiological delays, and other researchers have explicitly noted flowering extensions. For example, [52] working with mung bean reported that higher EMS levels prolonged the time to flowering, likely due to mutational interference with developmental pathways. It is also recorded by [53,54], [55] in black gram (*Vigna radiata*). In cowpea, [56] observed an earlier onset of flowering and a decrease in days to 50% flowering in an EMS-treated yardlong bean (*Vigna sesquipedalis*) population, accompanied by reduced maturity duration. Such wide variation in flowering time due to EMS has also been reported by [15], who found that their M□ cowpea population showed continuous distribution for days to flowering far beyond the range of the wild type. This implies that EMS created both later- and earlier-flowering individuals. Notably, [15] identified dozens of mutants that flowered significantly (P<0.05) earlier than the parent variety, indicating the emergence of new earliness traits. These results suggest that specific mutations can accelerate flowering, potentially by knocking out repressors of flowering even as overall vigor is reduced. In our mutants, while the predominant trend at high EMS dosages was delayed flowering and maturity, a few early-blooming mutants were observed at lower EMS treatments, indicating increased phenological variation. Our results confirm that EMS mutagenesis increased the variance in flowering time. While early flowering is advantageous for drought avoidance and multi-cropping systems, our evaluation of the EMS population did not yield any early-flowering mutants; instead, such occurrences were observed in the control group. This outcome aligns with results previously reported in mung bean and other pulse crops.

##### Effect of EMS on Days to First Harvest

The present study showed EMS treatment at all three concentrations produced a highly significant delay in days to first harvest (*dfh*) compared to the untreated control. No significant differences were detected among the three EMS concentrations themselves, again pointing to a threshold response wherein even the lowest concentration was sufficient to substantially delay the onset of first harvest, with no additive effect at higher doses. The delays in days to first harvest observed in the present study are a logical downstream consequence of the earlier-documented delays in days to first flower and days to 50% flowering and are consistent with the general expectation that any disruption of floral transition will propagate through the reproductive developmental timeline. In cowpea, pod development and seed fill are closely coupled to the timing of anthesis, and a delay of in flowering onset would be expected to produce a delay of similar harvestable pod maturation as observed here. [25], applied EMS at 10 mM for 6 hours to cowpea varieties *IT84E-124* and *Vita 7*, reported variants in maturity date among the spectrum of mutations recovered in the M□ generation, demonstrating that EMS can disturb the genetic determinants of cowpea maturation timing even at concentrations below those used in the present study.

In this study, the mutation frequencies recorded as 18.75%, 9.10%, and 12.50% for 20, 40, and 80 mM respectively, do not follow a simple linear dose-response pattern, with the lowest EMS concentration producing the highest proportion of phenotypic variants for this trait. This is consistent with the broader mutagenesis literature indicating that higher doses of chemical mutagens tend to produce proportionally more lethal or cryptic mutations rather than phenotypically viable and selectable variants. The dose of EMS concentration and exposure duration both affect the viability of induced mutations, and the frequency of viable mutations declined at concentrations that approach or exceed the LD□□, as higher doses shift the mutation spectrum increasingly toward deleterious rather than functional alleles [57].

A similar study by [58] who evaluated EMS effects on quantitative traits in garden pea (*Pisum sativum* L.) in the M□ generation found that variation due to EMS treatment was significant for days to maturity, with increasing EMS concentration generally associated with delayed maturity, and noted that such delays in maturity were consistent across concentrations in a manner similar to flowering time responses. The strong similarity between *dfh* results across 20–80 mM in the present cowpea study, and the absence of a further dose-dependent increment beyond the initial shift from the control, mirrors this threshold pattern observed in garden pea. The relatively higher mutation frequency at 20 mM for days to first harvest therefore suggests that sub-lethal doses may be more productive for generating harvestable maturity variants in cowpea, though this interpretation must be qualified by the relatively small population sizes in this M□ study.

##### Effects of EMS on Days to 50% Mature Pods

Coefficients of variation were low and comparable across all treatments (9.1–10.5%), indicating that EMS did not increase population-level variance in pod maturation timing, but instead produced a coherent directional shift in mean maturity across the mutagenized populations. Mutation frequencies of 39.58%, 25.45%, and 23.21% for 20, 40, and 80 mM respectively again followed the declining trend with increasing EMS dose observed for *dfh,* reinforcing the interpretation that the 20 mM dose generated the highest proportion of viable phenotypic variants for maturation-related traits. The delay in *d50mp* closely parallelled the delay in *dfh* across all treatments, with the absolute number of days between first harvest and 50% pod maturity remaining stable across the control (approximately 30 days) and EMS treatments (approximately 28–30 days). This similarity suggests that EMS disrupted the upstream developmental program governing the onset of reproductive maturation, most plausibly through mutations affecting the floral transition pathway, rather than the rate of pod filling or seed development intrinsically. A study on EMS mutagenesis of fodder cowpea (var. *Aiswarya*) by [59] reported that days to maturity was among the quantitative traits showing significant variation in the mutated population in M□, with EMS-induced mutants covering a broad range of maturity classes including both early and late maturing individuals relative to the control[44]. The observation of delayed maturity in the present M□ study without evidence of any early-maturing variants at the population mean level is consistent with the expectation that dominant or co-dominant mutations, which are the only classes detectable at the phenotypic level in M□, are more likely to disrupt rather than accelerate developmental programs. Stable phenotypic variation in both directions, including earlier maturity, is expected to be more fully expressed from M□ onwards, as the first non-chimeric generation in which homozygous recessive mutations can be phenotypically detected [13,60].

The agronomic implications of delayed maturity in EMS-mutagenized cowpea M□ populations deserve consideration. While late maturity is undesirable from a production standpoint in environments with short growing seasons, it may represent a useful source of variation for adaptation to longer-season cropping systems or for intercropping arrangements where extended vegetative and reproductive periods are advantageous. The importance of early maturing mutations for short-season environments has long been recognized in cowpea improvement programs, with early maturing selections serving as candidates for areas with limited rainfall and as relay crops in rice paddies. In contrast, the late-maturing phenotype observed in the current M□ population may harbor individuals carrying alleles that, in heterozygous combination, mask early maturing recessive mutations, suggesting that early maturity mutants with direct breeding utility may emerge from the M□ onwards upon selfing and screening [61].

The results for *dfh* and *d50mp* demonstrated that EMS mutagenesis at concentrations of 20–80 mM induces a coherent and significant delay of 9–11 days in both harvest maturity and pod maturation in cowpea under M□ conditions. This delay is may be attributed to the dominant expression of EMS-induced mutations disrupting the genetic flow of floral transition and reproductive development, consistent with findings in cowpea and other legumes. The absence of dose-dependent differentiation among EMS treatments suggested a threshold effect for maturity traits, while declining mutation frequencies at higher doses indicate that 20 mM may represent the most productive concentration for generating viable phenotypic variants in maturity-related traits. Evaluation of M□ and subsequent generations will be essential to characterize the full spectrum of maturity variation induced, including the recovery of potentially valuable early-maturing mutant classes.

#### Effects of EMS Mutagenesis on Yield-related and Yield traits

##### Non-Significant Traits: Number of Pods per Peduncle, Number of Pods per Plant at Maturity, Number of Seeds per Pod, Number of Seeds per Plant, Pod Width, and Percent Seed Abortion

EMS treatment at all three concentrations (20, 40, and 80 mM) produced no statistically significant differences from the control in number of pods per peduncle (*npp*), number of pods per plant at maturity (*nptm*), number of seeds per pod (*nsp*), number of seeds per plant (*nspt*), pod width (*pdw*), or percent seed abortion (*psa*). The lack of significant and notable mutants in the M□ generation is likely due to these traits being highly polygenic and developmentally buffered, which restricts observable changes [4,62–64]. These findings suggest that a single generation of EMS exposure at these doses were insufficient to consistently disrupt the genetic architecture underlying these yield components in the M□ generation. However, significant mutants were identified for these traits in M□ populations, where recessive mutations are revealed, and chimerism is resolved [15,65-Click or tap here to enter text.67]. Moreover, higher EMS doses often reduce pod and seed numbers due to partial sterility or lower pollen viability, as seen in legumes [4,34,53].

[13] reported wide distribution of yield-related traits including number of pods per plant, number of seeds per pod, and number of seeds per plant primarily in the M□ population rather than M□, noting that the frequency of viable mutations depends strongly on treatment conditions and that functional mutations induced by EMS through point mutation have a greater chance of being dominant or co-dominant with only those classes easily detected by phenotypic observation in the M□ generation[13]. The absence of significant mean shifts in these traits in the current M□ study therefore may reflects the recessive nature of many yield-component mutations rather than an absence of induced mutations.

[31] reported highly significant mutants with improved performance for number of pods per plant, number of seeds per pod, number of locules per pod, and pod length were identified in EMS-mutagenized cowpea populations, but their detection required evaluation of the M□ generation, reinforcing the need to advance mutagenized material to subsequent generations for meaningful yield screening. Similarly, [25] reported increases in number of seeds per pod, peduncles per plant, and 100-seed weight in advanced mutant lines of cowpea varieties ***IT84E-124*** and ***Vita 7*** following mutagenesis with EMS at 10 mM and NaN□, though these improvements were characterized from M□ onwards rather than M□ plants [59]. The high coefficients of variation recorded in this study for *nsp*, *nptm*, *nspt*, and *psa* across all treatments, including the control, indicated that these traits are inherently variable and may require larger sample sizes or later generation evaluation to detect treatment effects. The notably high CV for *psa* reflected the expected wide individual variation in seed set across a mutagenized population. The higher performance in other yield traits exhibited by mutants may be attributed to pleiotropy, whereby mutation in one trait affects others [61], which further complicates detection of significant mean shifts in individual yield components when evaluation is confined to M□.

##### Effects of EMS on Pod Length

Pod length (*pdl*) showed a significant overall treatment effect, with a dose-dependent reduction across EMS concentrations. Pairwise comparison confirmed a significant difference only between the control and the 80 mM treatment, this suggest that the reduction in pod length became statistically detectable only at the highest EMS concentration. The highest mutation frequency for *pdl* of 21.43% was observed in 80-mM treatment, which indicated that the 80 mM dose induced the greatest proportion of detectable variants, consistent with a dose-response relationship in this trait. The reduction in pod length with increasing EMS concentration is consistent with findings in the broader mutagenesis literature for legumes. Studies on fodder cowpea in the M□ generation reported reductions in several quantitative characters including pod length with increasing EMS concentrations, with reductions in quantitative traits observed to be colinear with increased mutagen dose irrespective of the mutagen used [59]. This pattern has been attributed to the mutagenic disruption of genes governing cell elongation and pod development [26]. In legumes, the activity of the inflorescence meristem determines the number and size of flowers and pods at each node through the growth of the floral meristem, and mutations affecting this developmental pathway can alter pod size and architecture [26]. EMS-induced point mutations at loci controlling pod wall cell division or expansion would therefore be expected to reduce pod length in a dose-dependent manner [68], as observed here.

In contrast, some advanced-generation mutagenesis studies have reported enhanced pod length in selected mutant lines. [69],obtained aphid-resistant cowpea mutants from gamma-irradiated populations with increased pod length among their improved attributes, while the same study noted that lines with significant increases in yield parameters were selected from M□ screening [25]. The apparent contradiction between M□ reductions and M□ improvements in pod length likely reflects the phenomenon of heterozygous M□ plants masking recessive gain-of-function alleles, which only become homozygously expressed and selectable in subsequent generations [70]

##### Effects of EMS on Number of Locules per Pod

Number of locules per pod (*ln*) showed the most pronounced and dose-dependent response to EMS treatment among all traits evaluated. All pairwise comparisons among treatments were statistically significant, and the mutation frequency increased with concentration, which reached 50% at 80 mM. The CV also increased with EMS dose, which reflected the expanding phenotypic diversity generated at higher concentrations. The strong dose-dependent reduction in locule number is particularly notable from a breeding standpoint, as locule number is a primary determinant of the number of seeds per pod and therefore of pod yield potential. In legumes, the identity and activity of the inflorescence meristem governs flower and pod architecture, and mutations in genes regulating meristematic activity can directly alter the number of seeds per pod and overall pod structure[68].

The significant reduction in *ln* at 40 and 80 mM, without a corresponding significant reduction in *nsp*, suggests that while EMS disrupted the developmental program responsible for locule number, the individual seed development process within remaining locules were largely maintained. This dissociation between locule number and seeds per pod may indicate that some locule positions failed to initiate rather than that initiated locules failed to complete seed set. [31] noted that highly significant mutants with improved performance for number of locules per pod were identified in EMS-mutagenized cowpea in M□ populations, demonstrating that both directional increases and decreases in locule number are achievable through EMS mutagenesis depending on the specific mutation event. The predominance of locule-reducing variants in the current M□ population is consistent with the expectation that dominant loss-of-function mutations more likely to be expressed in heterozygous M□ plants tend to disrupt rather than enhance developmental programs. Alleles conferring additional locules, which would likely require gain-of-function or additive effects, may be present in the population but will only become phenotypically apparent upon homozygosity in M□.

##### Effects of EMS on Seed Dimensions — Length, Width, and Thickness

The three seed dimension traits; seed length (*sdl*), seed width (*sdw*) and seed thickness (*sdt*) collectively exhibited highly significant responses to EMS treatment, but with notably different dose-response patterns across the three concentrations. ***Seed length*** increased with EMS treatment, as all comparisons between the control and treated groups were highly significant. However, there was no significant difference between the 40- and 80-mM treatments, indicating a plateau above 40 mM. The coefficient of variation decreased with increasing EMS dose, showing reduced variability. This suggested that EMS caused a consistent shift toward longer seeds rather than increasing variation. This pattern was consistent with the expression of mutations that promoted seed elongation, leading to a uniform directional effect across the population.

***Seed width*** followed a consistent but shallower pattern of EMS-induced increase. All three EMS concentrations were significantly wider than the control, though none differed significantly from one another indicating again a threshold effect where even the lowest dose captured the full directional shift achievable for this trait. The absence of dose-dependent differentiation among EMS treatments for seed width, combined with the significant and consistent departure from the control, points to a single effective threshold below 20 mM at which EMS disrupted or altered the genetic regulation of seed width, with no further phenotypic increment at higher concentrations.

**Seed thickness**, by contrast, produced the most complex and non-linear response of the three dimensions. The 20 mM treatment was the only concentration to produce a significant increase over the control, while both the 40 mM and 80 mM treatments were statistically indistinguishable from the control but significantly lower than the 20 mM treatment. The mutation frequencies for *sdt*, 12.50%, 1.82%, and 1.79% at 20, 40, and 80 mM respectively showed by far the steepest drop between the lowest and intermediate doses of any trait in this study, suggesting that the genetic targets for seed thickness are highly sensitive to EMS dose and that concentrations above 20 mM largely neutralized or reversed the thickness-enhancing mutations, possibly by simultaneously inducing compensatory or antagonistic mutations in related seed developmental pathways.

The divergence in response among the three seed dimensions is agronomically and biologically meaningful. Seed length responded with a strong, progressive, dose-dependent increase that plateaued above 40 mM; seed width responded with a uniform, threshold-type increase independent of dose; and seed thickness showed a transient, concentration-restricted increase that was not maintained above 20 mM. This dissociation among the three axes of seed growth implies that seed length, width, and thickness in cowpea are at least partially under distinct genetic control, and that EMS mutagenesis differentially targeted these control systems across the tested concentration range. Consistent with this interpretation, QTL mapping studies in cowpea have identified distinct genomic regions controlling pod and seed dimensions, including pod length and 100-seed weight, suggesting that seed size traits can be genetically decoupled and are governed by a combination of shared and independent loci [71]. The increases in seed length and width documented here are consistent with findings from induced mutagenesis studies in cowpea and related legumes. [25] reported significant increases in seed weight among EMS-mutagenized cowpea lines, with traits including plant height, seed weight, and protein content increasing in mutant populations compared to untreated controls. The outcomes were attributed to EMS altering structural gene loci governing these quantitative traits[25]. [72] similarly documented substantial increases in 100-seed weight in gamma-irradiated cowpea mutants, with 100-seed weight rising from *16.95 g* in the control to *22.80 g* in selected M□ mutant lines, and observed that seed size traits displayed high heritability, indicating a strong genetic basis amenable to selection.

The pattern across all three seed dimensions, thus progressive elongation in length, uniform widening, and transient thickening points toward a net increase in overall seed volume in EMS-treated populations, predominantly driven by enhanced length and, to a lesser degree, width. This three-dimensional enlargement is likely to translate into increased individual seed weight, which is consistent with the significant increases in seed weight (*swg*) documented in this study. [13]attributed high performance in multiple yield traits exhibited by EMS-mutagenized cowpea mutants partly to pleiotropy, whereby mutation in one gene affects several related developmental traits. This mechanism may explain the coordinated, if not perfectly similar, increases across seed length, width, weight, and, at 20 mM, thickness observed in the current M□ population. As with all M□ data, the observed means represented population-level responses from heterozygous individuals and should be interpreted cautiously as indicators of induced genetic variation rather than fixed trait values.

Stable phenotypic expression of EMS-induced seed dimension changes is expected from M□ onwards, as the first non-chimeric generation in which homozygous alleles become expressed and individually selectable [13]. Selection in M□ for individuals combining large seed length, adequate width, and maintained thickness particularly among progeny of 20 and 40 mM-treated plants where mutation frequencies for seed dimension traits were highest, offers the most promising route to isolating fixed large-seeded mutant lines of breeding value.

##### Effects of EMS on Seed Weight

Seed weight (*swg*) showed the most pronounced and consistently positive response to EMS treatment among all yield traits evaluated. All three EMS concentrations produced significantly heavier seeds than the control. Pairwise comparisons confirmed that 20 and 40 mM did not differ significantly from each other, but both were significantly heavier than 80 mM, which in turn remained significantly heavier than the control. This pattern describes a partial dose-dependent attenuation: maximum seed weight enhancement occurred at 20–40 mM, with a decline though still significant improvement over the control at 80 mM. Mutation frequencies of 54.17%, 49.10%, and 16.07% at 20, 40, and 80 mM respectively confirm the well-documented inverse relationship between EMS dose and the proportion of viable, selectable phenotypic variants, as higher concentrations shift the mutation spectrum increasingly toward lethal or cryptic alleles rather than productive, phenotypically expressed changes[13]. The significant increase in seed weight across all EMS treatments is consistent with findings in cowpea and related grain legumes subjected to chemical and physical mutagenesis. [25], applying EMS at 10 mM for 6 hours to cowpea varieties ***IT84E-124*** and ***Vita 7***, reported significant increases in seed weight and other yield parameters among selected mutant lines, attributing improvements in polygenic characters like yield to changes in simply inherited traits or mutations at structural loci. [72] similarly reported significant increments in 100-seed weight in gamma-irradiated cowpea mutants at M□ and M□ generations, observing increases from *16.95 g* in the control to *22.80 g* in selected mutant lines, a proportional gain broadly comparable to that recorded in the 20- and 40-mM treatments of the current study.

The comparable increase in seed weight with the previously documented increases in seed length and seed width supports an interpretation of coordinated enhancement of seed size across multiple dimensions at low-to-intermediate EMS concentrations. [59], who used EMS to develop improved fodder cowpea populations, reported significant variation in seed yield per plant in M□ populations, noting that mutations altering seed morphology traits tend to produce coordinated changes in associated yield components, consistent with pleiotropic gene action or tight genetic linkage among seed size and weight loci. The reduction in seed weight at 80 mM, while still above the control, is likely attributable to higher mutagenic load at this concentration interfering with genes governing seed filling efficiency, endosperm development, or photosynthate partitioning effects that would be expected to partially offset the seed-enlarging mutations still being expressed from this dose.

##### Effects of EMS on Yield Per Plant

Yield per plant (*yld*) showed a significant overall treatment effect, but with a more limited and dose-restricted pattern than seed weight. Only the 20 mM treatment produced a statistically significant increase in yield relative to the control (*61.19 ± 3.34 g* vs. *48.79 ± 3.01 g*), representing an approximate 25% yield advantage. The 40 mM (*58.02 ± 2.59 g*) and 80 mM (*51.16 ± 2.87 g*) treatments produced numerically higher yields than the control but were not significantly different from it, nor from the 20 mM treatment. The high coefficients of variation across all groups (33.1–42.0%) reflect the inherently variable nature of yield per plant in M□ populations, a consequence of the heterogeneous mutational load across individuals and indicate that a portion of the variance attributable to EMS effects may have been obscured by this within-treatment variability. Mutation frequencies of 16.67%, 12.72%, and 10.71% for 20, 40, and 80 mM respectively were low and declined modestly with increasing EMS concentration, consistent with the modest treatment effect sizes recorded for this trait.

The significant yield increase at 20 mM is agronomically important, even within the context of M□ limitations. [59] documented significant variation in seed yield per plant in EMS-mutagenized cowpea M□ populations and concluded that EMS mutagenesis is effective in inducing yield-relevant genetic changes, and that seed yield and days to maturity were among the quantitative traits showing significant variation in the mutated population. The pattern observed in the current study which showed significant yield improvement at the lowest EMS concentration but non-significant trends at higher doses is consistent with the principle that sub-lethal mutagen doses are most productive for generating viable, yield-enhancing alleles. [13] attributed high yield performance in EMS-mutagenized cowpea mutants to EMS altering genes responsible for this trait, and additionally to pleiotropy, whereby mutation in one trait affects others a mechanism that is particularly relevant here, as the yield increases at 20 mM likely reflect the combined downstream effects of increased seed weight, enhanced seed dimensions, and maintained seed number documented in the same treatment group.

The overall yield results, viewed alongside the seed weight and seed dimension data, present an internally consistent picture: 20 mM EMS induced the broadest suite of positive seed quality and yield mutations in this M□ population, while 40 and 80 mM produced proportionally more neutral or disruptive mutations that dampened the yield response. As with all M□ observations, however, these means represent population averages across heterozygous individuals and cannot directly predict the yield potential of specific mutant lines. The identification and selection of superior individual mutant plants rather than evaluation of population means will be the critical next step, as demonstrated by, [72] who identified twelve putative cowpea mutants combining early maturity with significantly higher seed yields than the parental control through individual plant selection from M□ and M□ generations. Advancing the current population to M□ and screening individual lines for the convergence of high seed weight, appropriate seed dimensions, and elevated yield per plant represents the most productive strategy for exploiting the genetic variation induced by EMS in this study.

In our research, high-yielding mutants, such as B09, typically produce larger seeds with higher seed weight, a favorable combination for yield improvement. Similarly, [25] and [72] reported cowpea mutants with enlarged seeds and improved yield performance, consistent with the superior lines identified in the present study. Yield stimulation following mutagenic seed treatments has also been documented in other legumes. Increased yield or yield components after mutagen exposure have been reported in pigeonpea [73], mung bean [74], and urdbean treated with gamma rays, EMS, and sodium azide [75]. These studies collectively support the notion that induced mutagenesis can unlock favorable allelic variation controlling yield traits.

In contrast, many mutants, particularly from the highest EMS dose (80 mM), exhibited reductions in yield and its components, including pod length, number of locules, seed size, and seed weight. Such adverse effects of high mutagen doses are well documented. [76] In okra and [77] in petunia, it was reported that mutagenesis frequently produces lines with reduced reproductive output. In pigeon pea (*Cajanus cajan*), [78] observed that EMS doses beyond the optimal range led to sharp declines in pollen fertility and seed yield. At the same time, [79] reported positive yield shifts at lower mutagen doses and negative shifts at higher doses. Our results closely mirror this dose-dependent pattern. Yield depression at elevated doses of irradiation or chemical mutagens has been widely reported. Similar declines were observed by [52] in *Gloriosa superba*, [53] in *Vigna radiata* under EMS and DES treatments, [80] in *Vigna mungo*, and [81] in *Vigna radiata*. These findings reflect the accumulation of deleterious mutations at higher mutagen doses, which can impair physiological and reproductive processes. Conversely, [4] reported stimulation of seed weight in pigeon pea under combined mutagenic treatments, suggesting that moderate doses may enhance specific yield components. Similar observations were reported by [82], [83], and [26] across various pulse crops, reinforcing the dose- and trait-dependent nature of mutagenic responses.

One elite mutant (B33) from the 20 mM EMS treatment in the current study produced approximately 25% higher seed yield than the control, accompanied by a modest increase in 100-seed weight. This indicates that EMS successfully generated yield-enhancing alleles. Similarly, [59] reported EMS-derived cowpea lines with significantly increased pod numbers confirm the potential of chemical mutagenesis to improve yield. It is also noteworthy that the frequency of desirable mutants was low, consistent with previous reports. [15] estimated that only 0.1–1% of mutants exhibited significantly improved yield-related and yield traits. Similarly, although a few promising mutants were identified in the present study, the majority performed at or below the level of the control. This underscores the need to screen large mutant populations to capture rare beneficial events.

### Multivariate Analysis: PCA Biplot and Cluster Grouping

#### PCA Biplot of phenological, yield-related and yield traits

##### Variance structure and principal component interpretation

Principal component analysis was applied to the full set of 18 phenological and yield-related traits across 190 M□ cowpea genotypes to characterize the multivariate structure of EMS-induced variation. The first eight principal components collectively explained 76.2% of the total phenotypic variance, with PC1 and PC2 accounting for 17.7% and 15.1% respectively a combined 32.8% of variance captured in two dimensions. The relatively distributed variance structure across eight components reflects the complexity of the trait matrix, the phenotypically heterogeneous nature of M□ mutagenized populations, and the partial independence of phenological, reproductive, and seed morphology trait clusters. [85], evaluated cowpea breeding lines using multivariate analysis, similarly found that the first three principal components explained 76% of total variation and noted that groupings in biplots conformed well with dendrogram results, demonstrating that PCA effectively captures the major sources of agronomic differentiation in cowpea populations [7]. PC1 carried strong positive loadings from seed length (*sdl* = 0.73), seed width (*sdw* = 0.70), days to first harvest (*dfh* = 0.68), days to first flower (*dff* = 0.61), days to 50% mature pods (*d50mp* = 0.59), days to 50% flowering (d50f = 0.57), and seed weight (*swg* = 0.53). This axis therefore captures a composite gradient of phenological delay and seed dimensional enlargement, two trait groups that co-vary positively across EMS treatments in this present study, this reflects the tendency of mutagenized lines to simultaneously delay reproductive development and produce morphologically larger seeds. PC2 was dominated by highly positive loadings from number of seeds per plant (*nspt* = 0.96), yield per plant (*yld* = 0.95), and number of seeds per pod (*nsp* = 0.75), alongside negative loadings from number of pods per plant at maturity (nptm = −0.66), days to first flower (*dff* = −0.64), and days to 50% flowering (*d50f* = −0.68). PC2 thus represented a yield productivity axis, where high positive scores reflect superior reproductive output in terms of seed number and total yield, decoupled from phenological timing. Together, PC1 and PC2 defined a two-dimensional space within which the fundamental EMS-induced trade-off between phenological disruption, seed enlargement, and reproductive yield can be visualized at the individual genotype level.

##### Treatment group separation in the biplot

The biplot distribution of genotype scores revealed a clear dose-dependent spatial separation among EMS treatment groups. Control genotypes (A01–A31, 0 mM) formed a compact cluster on the extreme negative side of PC1, consistent with their early phenology, uniform developmental timing, and moderate to high yield performance relative to the mutagenized population. This clustering confirmed the low phenotypic variance expected of untreated plants within a uniform cultivar background and validates the internal coherence of the control group. [86], applied PCA to M□ Tswana cowpea mutant lines and similarly found that the control variety formed a separate cluster that diverged from all mutant lines in multivariate space, confirming that induced mutagenesis effectively altered the genetic makeup of the treated population in ways detectable by multi-trait analysis.

Genotypes from the 20 mM treatment (B01–B48) showed partial overlap with the control group but were moderately displaced toward the positive PC1 region and exhibited greater within-group dispersion, particularly along PC2. This intermediate positioning reflects the milder and more heterogeneous mutational load associated with the lowest EMS concentration were consistent with individual-level variation in mutation induction and expression at sub-lethal doses. Genotypes from the 40 mM (C01–C55) and 80 mM (D01–D56) treatments were progressively displaced toward the positive PC1 axis and displayed substantially greater scatter across both PC1 and PC2, indicating dose-dependent increases in phenological delay alongside widened variation in reproductive output. Multivariate PCA of cowpea germplasm has consistently shown that high phenotypic diversity, reflected by wider scatter in biplot space, creates greater opportunity for selection of superior genotypes, as variation among traits provides the basis for discriminating among individuals for multiple desirable characteristics simultaneously [87].

##### Trait vector interpretation and EMS-induced trade-offs

The present study showed the orientation of trait vectors in the biplot provided insight into the relationships among traits and the nature of EMS-induced phenotypic trade-offs. Vectors for the four phenological traits (*dff*, *d50f*, *dfh*, *d50mp*) were oriented horizontally towards the positive PC1 axis, this confirmed their strong association with the right-side positioning of high-dose EMS genotypes. The *sdl* and *sdw* vectors were directed toward the lower-right quadrant, positive PC1 and negative PC2 which indicated that seed dimensional enlargement was associated with phenological delay but inversely related to yield productivity. In contrast, the *yld, nspt*, and *nsp* vectors pointed strongly toward the upper region of the biplot (positive PC2), which formed an approximately perpendicular angle to the *sdl*/*sdw* vectors. This near-orthogonal relationship between yield and seed size vectors quantitatively confirms the resource allocation trade-off implied by the individual trait analyses: EMS-induced genotypes that developed larger seeds tended to produce fewer seeds per pod and per plant, resulting in reduced total yield. PCA-driven multivariate analyses in other legume systems have similarly identified trade-offs between plant architectural and yield productivity traits loading on different principal components, where biplots revealed that selection based on single traits may inadvertently compromise performance in correlated traits and reinforced the need for multi-trait selection indices[88].

The *swg* vector was oriented toward the positive PC1 and moderately positive PC2 region which indicated that heavier seeds were associated with both phenological delay and, to a degree, maintained or enhanced yield performance, a positioning distinct from *sdl* and *sdw*, which loaded more strongly on the negative PC2 side. This suggested that seed weight integration partly captured the influence of both seed size and seed number components, reflecting its composite nature as a product of seed dimensions and density.

The *pdl* and *ln* vectors were directed upward left, with *pdl* loading strongly on PC2 and *ln* on both PC1 and PC2. These vectors were oriented broadly toward the central and upper biplot region, indicating that genotypes with longer pods and higher locule numbers were not confined to a single EMS treatment group and could be found across the treatment spectrum consistent with the individual trait analyses which showed that pod length was significantly reduced only at 80 mM while locule number showed a progressive dose-dependent reduction.

##### Identification of putative mutant classes

The biplot facilitated the graphical identification of three distinct classes of agronomically relevant putative mutants, each positioned in a different region of the two-dimensional space. High-yielding putative mutants were identified from genotypes clustered in the upper region of the biplot with positive PC2 scores, in close alignment with the *yld*, *nsp*, *nspt*, and *swg* vectors. These included B33, B28, B22, B31, D56, D45, D25, C5, C41, B26, and B21. The presence of genotypes from both 20 mM (B-series) and 40–80 mM (C and D series) treatments among this group indicated that yield-enhancing mutations were not exclusively associated with the lowest EMS dose, though most B-series genotypes in this cluster is consistent with the higher mutation frequencies and significant mean yield increase documented at 20 mM in the individual trait analyses. [86] similarly identified mutant lines with improved grain yield through PCA biplot analysis and noted that the distribution of mutagenized populations in biplot space, this reflected the interaction of genotype and mutagen dose that generated significant quantitative trait variation that could be exploited through targeted selection [86].

Large-seed putative mutants were identified from genotypes located along the negative PC2 axis in the direction of the *sdl* and *sdw* vectors, including D23, D7, D34, D47, C37, C39, C40, C42, B18, B11, and B8. These genotypes, from the 40- and 80-mM treatments, produced larger seeds but fewer in number, which resulted in lower overall yield. This pattern suggested that resources were redirected to produce fewer but larger seeds. Although this was not ideal for total yield, these genotypes were useful for breeding large-seeded varieties preferred by consumers and for studying the genetic basis of seed development in cowpea.

Early-phenology putative mutants were identified from the extreme negative side of PC1, opposite to the late-flowering and maturity vectors, and included B19 and C2 notably from the 20- and 40-mM treatments respectively. These genotypes showed accelerated phenological development relative to the control group, suggesting the presence of rare dominant or semi-dominant early-flowering mutations within the mutagenized population. Their identification is agronomically significant: previous studies in cowpea mutagenesis have confirmed that EMS causes mutations in genes responsible for maturity-related traits, and early maturing mutants are particularly valuable for short-season production systems and as relay crops in rice paddies where a compressed growing season is critical[13].

Genotypes exhibiting extreme positive PC1 scores with weak alignment to yield vectors including B9, B7, C46, and D20 were classified as severely delayed and low performing. These lines, positioned in the far-right region of the biplot with little positive PC2 displacement, represented the most mutagenized damaged individuals in the population, they likely carried multiple loss-of-function mutations with pleiotropic negative effects on reproductive development and yield. While unsuitable for direct advancement, they were retained as extreme phenotypes for genetic and physiological characterization, and as potential donors in targeted crosses where specific mutant alleles may segregate independently from the broader phenotypic impairment.

##### Broader utility of PCA in EMS mutation studies

In this study we reduced the dimensionality of the 18-trait dataset into two interpretable axes, the biplot enabled simultaneous visualization of dose-treatment group separation, inter-trait relationships, and individual genotype positioning relative to trait vectors all within a single analytical framework. This multi-trait perspective is particularly important in M□ studies where high within-treatment variance, heterozygosity, and chimerism create complex phenotypic distributions that are difficult to interpret through single-trait analyses alone. PCA biplot analysis of cowpea populations has been shown to effectively identify which traits account for the most variation among genotypes, allowing breeders to focus selection on the most discriminated trait combinations and to identify individuals that deviate meaningfully from the population center in the direction of target trait improvement. [89]. Also, [90], used multi-trait analysis to identify superior M□ individuals and reported that mutants with extremely high values in key traits, such as seeds per plant, could be distinguished from the population distribution. In the present study, the biplot confirmed that yield-related traits (*npp*, *nsp* and *swg*) and seed dimension traits loaded most strongly on the first two principal components, this provided the clearest discrimination among EMS-treated lines, while traits with minimal biplot contribution such as *psa* and *pdw* had limited power to differentiate among genotypes at the multivariate level.

The clear spatial separation between high- and low-yielding mutants, and between early- and late-phenology groups, demonstrated that the genetic variation induced by EMS at 20–80 mM was sufficient to generate agronomically meaningful divergence, detectable at the population level in the M□ generation. Confirmation of the most promising putative mutants particularly the high-yielding group (B33, B28, B22, B31, D56, D45, D25, C5, C41, B26, B21) and the early-phenology group (B19, C2) through M□ progeny testing and subsequent generation advancement remains essential before any breeding value can be assigned.

#### Hierarchical Cluster Analysis of EMS-Mutagenized Cowpea

Hierarchical cluster analysis using Ward’s linkage method effectively partitioned the 190 M□ cowpea genotypes into six phenotypically distinct clusters, each characterized by a unique combination of phenological, yield-related and yield trait profiles. The resolution of six clusters from a single mutagenized population is consistent with reports from other induced mutagenesis studies in cowpea and related legumes, which have commonly observed between three and six discrete phenotypic groupings following mutagen treatment, reflecting the breadth of the genetic variation induced [86,91]. The clustering outcome in the present study demonstrated that EMS mutagenesis at 20–80 mM generated sufficient phenotypic divergence across 18 traits to resolve multiple distinct genotypic classes within a single cultivar background, a result that underscores the utility of Ward’s method for structuring genotype selection in M□ mutation populations.

##### Cluster 1: high-yielding EMS-derived genotypes

Cluster 1, characterized by prolonged phenological development and superior yield performance, contained disproportionately high proportions of the 20- and 40-mM treatment groups, with 35.42% and 49.10% of these populations respectively assigned to this cluster. Cluster 1 contained the highest proportion of agronomically desirable genotypes, and most putative mutants selected for advancement. The combination of elevated seed weight (up to 28.68 g in D56), extended phenological development, and high yield per plant (up to 111.85 g in D56) in this cluster is consistent with reports of yield-enhancing mutations in EMS-mutagenized cowpea populations. [13] selected the top twenty mutants with high yield performance in terms of number of seeds per plant from their M□ EMS-mutagenized cowpea population, they reported that traits including higher number of pods per plant, increased protein content and high seed weights had been obtained through EMS mutagenesis in earlier cowpea studies. The genotypes identified in Cluster 1 in the present study include D56, D45, D25, B31, B28, B22, C41, and C5 exhibited yield values significantly exceeding the control range and represented the primary candidates for advancement to M□ and subsequent generation evaluation.

##### Cluster 2: control-dominated baseline cluster

Cluster 2, which represented the population’s baseline phenotypic profile and was dominated by control genotypes (61.29% of the 0 mM population), received minimal representation from EMS-treated lines. The concentration of control genotypes in Cluster 2 confirmed that the untreated population occupied a phenotypically distinct region of the multivariate trait space, separated from the majority of EMS-treated genotypes. The small number of EMS-treated lines assigned to this cluster notably C02 and C14, both from the 40 mM treatment, with early days to first harvest values of 38 and 39 days respectively, represented a class of putative early-phenology mutants that maintained control-like performance across most traits while showing accelerated crop development. [91], who developed new cowpea mutant genotypes through gamma irradiation, also reported that cluster groupings could be attributed to the origin of genotypes and the degree of genetic alteration induced, with control varieties consistently forming separate clusters that diverged from mutagenized lines in multivariate space [92]. This confirming the effectiveness of mutagenesis in altering the genetic makeup of the treated population [92].

##### Cluster 3: severely impaired high-dose genotypes

In contrast, Cluster 3 was characterized by delayed phenology, reduced locule number, and low yield, and it was mainly composed of genotypes from the 80 mM treatment, with 28.57% of that group assigned to this cluster. This cluster showed the most severely impaired phenotypes in the population. Representative genotypes B7, B9, C46, and D20 showed marked negative deviations from the control range across multiple traits simultaneously and were classified as low-value mutants unsuitable for direct advancement. This cluster is analogous to the poor-performance groups consistently reported in high-dose mutagenesis studies. [93] noted that higher mutagen doses tend to produce populations with reduced fertility, delayed maturity, and inferior agronomic performance, and recommended against their use in breeding programmes where the objective is to recover viable, productive mutant lines[94]. While the genotypes in Cluster 3 were not selected for advancement, they were retained as phenotypic reference material, as the extreme mutations they carry may be informative for gene function studies or for identifying linkage between yield-impaired and yield-neutral alleles in subsequent molecular characterization.

##### Cluster 4: early-flowering high-yield mutants

Cluster 4 contained a distinctive combination of early days to first flower and days to 50% flowering with extended days to first harvest and elevated yield, a trait profile not represented in any other cluster. The presence of B19 (*yld* = 93.59 g, *dff* = 36 days) and B33 (*yld* = 125.44 g, *dff* = 37 days) in this cluster is particularly noteworthy: these genotypes achieved the highest individual yields in the entire population while retaining comparatively early flowering phenology, a combination of traits of direct relevance to short-season cowpea production. EMS-induced early maturing mutants have been specifically highlighted as valuable in cowpea breeding because of their ability to escape drought or tolerate insect damage due to their short reproductive phase, with early mutants with days to flowering of 38–41 days having been isolated from M□ EMS-mutagenized cowpea populations in earlier studies [13]. The presence of such genotypes in Cluster 4 across the 20- and 40-mM treatments, but not exclusively at the lowest dose, reinforces the value of evaluating mutant lines individually rather than making selection decisions based solely on treatment concentration.

##### Cluster 5: moderate performance with seed weight improvement

Cluster 5, composed exclusively of EMS-treated genotypes from all three treatment groups, was characterized by moderately delayed phenological traits, elevated seed weight, and reduced locule number, resulting in yield performance generally below the Cluster 1 and Cluster 4 groups but above the impaired lines of Cluster 3. The absence of control genotypes from this cluster confirms that the trait profile it represented, elevated seed weight combined with moderately delayed development and was not expressed within the untreated population and is therefore a product of mutagenesis. Genotypes in this cluster may still harbor breeding value as parents for crosses targeting seed quality improvement, even if their direct yield performance does not match the elite lines identified in Clusters 1 and 4.

##### Cluster 6: large-seeded, seed-weight-driven yield

Cluster 6 was the smallest and most phenotypically distinctive group in the analysis, contained only 9 genotypes from the 20- and 80-mM treatments. These genotypes combined large seed dimensions and high seed weight with moderate phenological delay and relatively low seed number per pod, producing elevated yield per plant primarily driven by individual seed weight rather than seed number. Selected genotypes from this cluster, including B07 (*swg* = 29.99 g, *yld* = 67.18 g) and D36 (*swg* = 28.29 g, *yld* = 74.69 g), represented a distinct agronomic type that may be particularly valuable for breeding programmes targeted at large-seeded, consumer-preferred market classes. [95], developed high-yielding cowpea mutant lines through gamma irradiation and sodium azide, similarly used dendrogram analysis to separate control and mutant plants into distinct clusters, confirmed that considerable genetic variability was generated and that genetically diverged mutants could be employed directly as mutant varieties or as parents in crossbreeding programmes for advanced improvement and trait fixation[95].

##### Broader utility of cluster analysis in EMS mutation studies

This dose-stratified clustering pattern is consistent with the well-established principle that lower and intermediate mutagen doses preferentially generate viable, selectable phenotypic variants, while higher doses shift the mutation spectrum toward lethal or disruptive alleles. [93] confirmed this pattern in cowpea, they demonstrated that lower and intermediate mutagen doses induced higher frequencies of morphological mutations with the least biological damage and recommended these doses for future breeding programmes to obtain a high spectrum of desirable mutants[94]. The dominance of 40 mM-derived genotypes in Cluster 1 and of 80 mM-derived genotypes in Cluster 3 precisely mirrors this dose–performance gradient. However, EMS dose was not a perfect predictor of cluster assignment and individual genotype performance must be considered on its own merits. Some 20 mM genotypes were assigned to lower-performing clusters, while some 80 mM genotypes clustered with better-performing lines. [13], in their EMS mutagenesis study of cowpea cv. *Asontem,* emphasized that the phenotypic expression of each mutant line is unique and determined by the specific gene(s) altered by the mutagen. [13] noted that the high yield performance of selected mutants may reflect EMS altering genes responsible for traits as well as pleiotropic effects whereby mutations in one trait affect related traits[13]. Results from the present study reinforced this interpretation: multivariate clustering based on actual trait performance provided a more reliable selection criterion than EMS dose alone.

The clustering outcomes confirmed that the elite putative mutants identified across Clusters 1, 4, and 6 shared a multi-trait profile that were not present in the control population, suggesting they carried functional alleles which favored yield, seed size, or early phenology that were induced by EMS treatment. Agronomic evaluation of cowpea mutant lines in Eswatini demonstrated that cluster analysis is a key non-parametric statistical technique that facilitates the identification of diverse genotypes with contrasting traits for breeding and production, and that genotypes grouped in high-yielding sub-clusters can be directly recommended for production or use as crossing parents in further improvement programmes[92]. The present study reinforces this conclusion, with the elite clusters identified here providing a structured and evidence-based shortlist of M□ genotypes for advancement to M□ and evaluation in replicated trials across environments.

### Conclusion

This study demonstrates the effectiveness of ethyl methane sulfonate (EMS) in inducing substantial genetic variability in cowpea (*Vigna unguiculata* (L.) Walp), thereby broadening the phenotypic spectrum for key agronomic, yield, and seed traits. Contrast analysis revealed significant treatment effects and highlighted superior mutant lines, while principal component analysis (PCA) identified yield and reproductive traits as the primary drivers of phenotypic variation. These findings were consistently supported by cluster analysis, which grouped mutants according to overall agronomic performance. The convergence of these analytical approaches strengthened the selection of putative mutants, including high-yielding (B33, B31, B28, B22, C41, C05, D25, D45, D56), early-phenology (B19), and large-seeded or prolific podding lines (D34, C39, C40).

As the study was confined to the M□ generation, further evaluation in the M□ and M□ generations is required to confirm trait stability, heritability, and breeding value. Complementary molecular analyses will be essential to explain the genetic basis of the observed variation and support marker-assisted selection. Overall, the selected mutants constitute valuable genetic resources for cowpea improvement and have strong potential to enhance productivity and global food security. Considering the documented genetic variability of cowpea for nutritional traits such as protein and mineral content, the EMS-induced mutants identified in this study may serve as valuable genetic resources for future breeding programs targeting both yield and nutritional quality improvement.

## Acknowledgments

HMK, RANK, IKA, and FO conceived and designed the study. HMK and RANK performed the experiments. HMK analyzed the data and drafted the manuscript. RANK and IKA provided critical revisions. All authors read and approved the final manuscript. We gratefully acknowledge the Department of Plant and Environmental Biology at the University of Ghana, Legon, for supporting this research.

## Supporting Information

S1 Table. Trait dataset and cluster grouping of individual M□ cowpea plants across EMS treatments and control.

S2 Table. PCA loadings and variance explained for traits in M□ cowpea.

