## Supplementary material for "Effect of Ethyl Methane Sulfonate Mutagenesis on Phenological, Yield-Related and Yield Traits in Cowpea *(Vigna unguiculata* (L.) Walp)": S1 Table

**S1 Table: Trait dataset and cluster grouping of individual $M_1$ cowpea plants across EMS treatments and control**

| Trt | ID | *dtg* | *dff* | *d50f* | *dfh* | *d50mp* | *pdl* | *ln* | *sdl* | *sdw* | *sdt* | *swg* | yld | Cluster group |
|---|---|---|---|---|---|---|---|---|---|---|---|---|---|---|
| 0mM | A01 | 4 | 40 | 60 | 67 | 89 | 12.94 | 13 | 8.94 | 5.6 | 4.3 | 23.2 | 55.68 | 1 |
| | A02 | 4 | 39 | 60 | 53 | 88 | 17 | 18 | 6.62 | 4.53 | 3.86 | 20.34 | 42.71 | 2 |
| | A03 | 3 | 40 | 56 | 49 | 87 | 13.24 | 21 | 8.98 | 5.4 | 4.13 | 17.94 | 43.05 | 2 |
| | A04 | 3 | 35 | 52 | 50 | 81 | 13.2 | 13 | 7.03 | 4.35 | 3.29 | 23.96 | 52.71 | 2 |
| | A05 | 3 | 43 | 66 | 58 | 93 | 14.24 | 18 | 7.22 | 4.7 | 4.02 | 20.08 | 43.38 | 2 |
| | A06 | 3 | 30 | 60 | 60 | 78 | 13.58 | 21 | 6.72 | 4.42 | 3.7 | 20.97 | 41.53 | 2 |
| | A07 | 4 | 40 | 55 | 47 | 75 | 13.8 | 12 | 9.26 | 5.71 | 4.19 | 24.87 | 34.82 | 3 |
| | A08 | 4 | 41 | 62 | 53 | 83 | 11.55 | 19 | 7.91 | 5.11 | 4.17 | 19.93 | 36.27 | 2 |
| | A09 | 4 | 37 | 52 | 41 | 67 | 15.4 | 14 | 6.16 | 4.44 | 4.41 | 20.92 | 72.81 | 2 |
| | A10 | 4 | 37 | 54 | 58 | 89 | 13.09 | 16 | 6.23 | 4.15 | 3.64 | 22.13 | 44.92 | 2 |
| | A11 | 4 | 40 | 62 | 45 | 74 | 15.29 | 18 | 7.37 | 4.84 | 4.29 | 21.1 | 31.66 | 2 |
| | A12 | 4 | 44 | 59 | 46 | 76 | 15.95 | 17 | 8.98 | 5.4 | 4.13 | 21.01 | 43.71 | 3 |
| | A13 | 3 | 43 | 60 | 49 | 87 | 17.58 | 16 | 6.99 | 4.46 | 3.81 | 17.82 | 42.78 | 2 |
| | A14 | 5 | 30 | 52 | 53 | 80 | 16.64 | 18 | 9.65 | 5.67 | 4.28 | 23.13 | 73.78 | 3 |
| | A15 | 4 | 39 | 54 | 47 | 80 | 15.57 | 17 | 8.24 | 5.25 | 4.13 | 19.75 | 40.09 | 3 |
| | A16 | 3 | 45 | 66 | 40 | 70 | 14.08 | 24 | 6.77 | 4.57 | 4.36 | 20.79 | 39.49 | 2 |
| | A17 | 4 | 31 | 54 | 43 | 66 | 17.21 | 12 | 6.74 | 4.34 | 3.42 | 20.36 | 61.08 | 2 |
| | A18 | 3 | 32 | 49 | 56 | 94 | 14.25 | 20 | 7.57 | 5.03 | 4.92 | 22.64 | 67.93 | 4 |
| | A19 | 2 | 35 | 59 | 42 | 72 | 15.3 | 13 | 6.21 | 4.46 | 4.43 | 20.61 | 54.41 | 2 |
| | A20 | 3 | 33 | 48 | 47 | 84 | 16.62 | 20 | 6.38 | 4.3 | 4.1 | 22.57 | 72.66 | 4 |
| | A21 | 4 | 40 | 57 | 44 | 71 | 13.33 | 17 | 6.43 | 4.47 | 4.19 | 18.36 | 50.5 | 2 |
| | A24 | 4 | 39 | 58 | 49 | 72 | 15.42 | 17 | 5.68 | 3.94 | 3.63 | 19.63 | 61.83 | 2 |
| | A25 | 3 | 39 | 59 | 50 | 70 | 12.83 | 21 | 6.24 | 4.19 | 3.71 | 24.67 | 51.81 | 2 |
| | A26 | 3 | 37 | 59 | 54 | 86 | 15.3 | 23 | 6.76 | 4.54 | 4.53 | 22.04 | 52.89 | 4 |
| | A27 | 4 | 32 | 53 | 53 | 84 | 16 | 16 | 11.59 | 6.08 | 4.82 | 19.21 | 22.86 | 3 |
| | A28 | 4 | 40 | 65 | 43 | 79 | 11.88 | 12 | 6.07 | 4.03 | 3.85 | 21.97 | 85.68 | 2 |
| | A29 | 2 | 39 | 57 | 44 | 81 | 14.92 | 17 | 8.49 | 5.13 | 3.92 | 23.24 | 23.24 | 3 |
| | A30 | 4 | 43 | 68 | 51 | 88 | 13.53 | 18 | 8.35 | 5.15 | 4.48 | 18.63 | 50.31 | 2 |
| | A31 | 3 | 40 | 64 | 40 | 69 | 14.9 | 13 | 6.61 | 4.53 | 4.52 | 18.39 | 74.66 | 2 |
| 20mM | B01 | 5 | 50 | 70 | 68 | 92 | 12.13 | 18 | 6.33 | 4.24 | 4.06 | 22.28 | 55.69 | 5 |
| | B02 | 4 | 46 | 66 | 59 | 98 | 13.23 | 21 | 6.65 | 4.38 | 3.66 | 26.08 | 28.69 | 5 |
| | B03 | 5 | 42 | 60 | 53 | 79 | 14.98 | 13 | 6.94 | 4.58 | 4.23 | 25.41 | 37.36 | 5 |
| | B04 | 6 | 43 | 60 | 60 | 100 | 16.28 | 18 | 9.26 | 5.71 | 4.19 | 25.92 | 54.44 | 1 |
| | B05 | 5 | 47 | 66 | 62 | 82 | 13.8 | 21 | 10.1 | 5.78 | 4.92 | 25.17 | 49.83 | 1 |
| | B06 | 5 | 51 | 75 | 54 | 76 | 15.75 | 12 | 10.35 | 5.76 | 4.91 | 25.9 | 62.15 | 6 |
| | B07 | 7 | 38 | 60 | 63 | 89 | 11.45 | 19 | 11.65 | 6.44 | 5.19 | 29.99 | 67.18 | 6 |
| | B08 | 5 | 49 | 74 | 68 | 102 | 15.25 | 14 | 9.85 | 5.67 | 5.12 | 22.68 | 41.27 | 1 |
| | B09 | 7 | 52 | 74 | 71 | 98 | 13 | 21 | 11.59 | 6.08 | 4.82 | 27.48 | 74.2 | 6 |
| | B10 | 6 | 47 | 63 | 56 | 81 | 14.31 | 23 | 7.03 | 4.48 | 3.79 | 20.06 | 46.14 | 2 |
| | B11 | 7 | 52 | 70 | 60 | 95 | 13.3 | 16 | 9.75 | 5.66 | 4.99 | 26.1 | 26.1 | 3 |
| | B12 | 3 | 41 | 59 | 49 | 87 | 17.7 | 13 | 10.4 | 5.9 | 5.16 | 26.32 | 48.7 | 6 |
| | B13 | 4 | 47 | 62 | 66 | 100 | 14.25 | 18 | 9.9 | 5.72 | 4.87 | 23.49 | 59.2 | 1 |
| | B14 | 7 | 53 | 69 | 58 | 87 | 14.5 | 21 | 7.18 | 4.88 | 4.76 | 24.86 | 45.75 | 5 |
| | B15 | 3 | 54 | 71 | 55 | 76 | 15.25 | 16 | 9.75 | 5.88 | 4.83 | 27.77 | 79.15 | 1 |
| | B16 | 5 | 53 | 75 | 64 | 102 | 14.7 | 18 | 9.75 | 5.68 | 4.85 | 26.24 | 75.56 | 1 |
| | B17 | 3 | 41 | 65 | 71 | 93 | 13.7 | 17 | 10.4 | 5.82 | 4.84 | 27.34 | 71.37 | 6 |
| | B18 | 6 | 54 | 71 | 62 | 98 | 15.5 | 12 | 10 | 5.62 | 4.89 | 28.33 | 45.32 | 3 |
| | B19 | 5 | 36 | 51 | 48 | 68 | 19.88 | 19 | 6.23 | 4.17 | 4 | 24.82 | 93.59 | 4 |
| | B20 | 5 | 34 | 57 | 52 | 72 | 14.7 | 14 | 6.77 | 4.57 | 4.36 | 26.63 | 42.6 | 5 |

Values highlighted red indicate trait means significantly (P<0.05) below the control, while green indicate values significantly (P<0.05) above the control.Key: Trt= Treatment ID=Genotype Identifier, dtg days to germination, dff = days to first flower, d50f = days to 50% flowering, dfh = days to first harvest, d50mp = days to 50% mature pods, pdl = pod length, pdw = pod width, ln = number of locules, nsp = number of seeds per pod, psa = percent seed abortion, sdl = seed length, sdw = seed width, sdt = seed thickness, swgt = seed weight, yld = yield per plant

**S1 Table: Trait dataset and cluster grouping of individual $M_1$ cowpea plants across EMS treatments and control (*cont'd*)**

| Trt | ID | *dtg* | *dff* | *d50f* | *dfh* | *d50mp* | *pdl* | *ln* | *sdl* | *sdw* | *sdt* | *swg* | *yld* | Cluster group |
|---|---|---|---|---|---|---|---|---|---|---|---|---|---|---|
| 20mM | B21 | 5 | 47 | 63 | 70 | 91 | 13.6 | 13 | 9.85 | 5.64 | 5.09 | 22.6 | 81.37 | 1 |
| | B22 | 6 | 36 | 56 | 70 | 97 | 12.1 | 20 | 7.04 | 4.98 | 4.85 | 21.91 | 88.96 | 1 |
| | B23 | 7 | 48 | 73 | 55 | 79 | 13.04 | 17 | 8.26 | 5.04 | 3.93 | 22.04 | 46.28 | 5 |
| | B24 | 6 | 50 | 74 | 52 | 75 | 13.35 | 13 | 8.7 | 5.13 | 4.91 | 17.81 | 18.7 | 3 |
| | B25 | 7 | 39 | 61 | 58 | 89 | 17.75 | 18 | 9.65 | 5.31 | 4.52 | 25.92 | 76.99 | 1 |
| | B26 | 4 | 47 | 72 | 64 | 100 | 14 | 21 | 10.1 | 5.54 | 4.87 | 24.26 | 91.45 | 1 |
| | B27 | 7 | 39 | 55 | 56 | 85 | 16.1 | 12 | 7.04 | 4.68 | 4.28 | 25.59 | 67.56 | 4 |
| | B28 | 4 | 39 | 60 | 67 | 87 | 12.13 | 19 | 7.3 | 4.75 | 4.04 | 23 | 103.5 | 1 |
| | B29 | 6 | 43 | 66 | 61 | 94 | 13.27 | 14 | 6.62 | 4.53 | 3.86 | 24.2 | 45.73 | 5 |
| | B30 | 7 | 40 | 61 | 63 | 95 | 14.1 | 13 | 6.46 | 4.29 | 4.1 | 23.82 | 28.58 | 5 |
| | B31 | 3 | 39 | 60 | 58 | 98 | 14.85 | 20 | 9.9 | 5.46 | 5 | 24.65 | 107.22 | 1 |
| | B32 | 6 | 47 | 71 | 49 | 85 | 14.22 | 17 | 6.14 | 4.06 | 3.87 | 22.04 | 66.11 | 4 |
| | B33 | 6 | 37 | 61 | 56 | 85 | 16.33 | 19 | 6.1 | 4.05 | 3.31 | 27.88 | 125.44 | 4 |
| | B34 | 7 | 41 | 57 | 53 | 85 | 12.44 | 17 | 7.04 | 4.98 | 4.85 | 22.2 | 38.86 | 5 |
| | B35 | 7 | 51 | 68 | 56 | 91 | 14.75 | 13 | 10.05 | 5.24 | 3.56 | 26.06 | 63.85 | 5 |
| | B36 | 4 | 33 | 54 | 64 | 100 | 14.1 | 19 | 10.15 | 5.18 | 3.91 | 23.62 | 64.48 | 1 |
| | B37 | 7 | 53 | 73 | 62 | 82 | 14.35 | 17 | 11.25 | 5.77 | 3.71 | 26.15 | 82.38 | 1 |
| | B38 | 5 | 43 | 61 | 61 | 81 | 14 | 13 | 10.3 | 5.01 | 3.84 | 27.88 | 42.94 | 5 |
| | B39 | 6 | 54 | 71 | 49 | 85 | 13.25 | 19 | 10.4 | 5.67 | 4.46 | 26.71 | 86.53 | 1 |
| | B40 | 7 | 37 | 58 | 61 | 90 | 16.6 | 15 | 10.2 | 5.5 | 4.4 | 26.06 | 68.8 | 6 |
| | B41 | 7 | 33 | 50 | 60 | 99 | 17.52 | 18 | 6.43 | 4.47 | 4.19 | 25.04 | 66.61 | 4 |
| | B42 | 5 | 49 | 74 | 69 | 107 | 12.7 | 20 | 6.74 | 4.34 | 3.42 | 26.21 | 46.13 | 5 |
| | B43 | 3 | 37 | 56 | 58 | 96 | 12.25 | 18 | 9.3 | 4.73 | 3.72 | 23.01 | 68.33 | 5 |
| | B44 | 6 | 48 | 70 | 69 | 107 | 14.43 | 19 | 6.65 | 4.66 | 4.27 | 21.38 | 29.94 | 5 |
| | B45 | 3 | 48 | 67 | 62 | 96 | 13.75 | 20 | 10.1 | 5.57 | 4.08 | 23.17 | 76.45 | 1 |
| | B46 | 6 | 46 | 65 | 55 | 80 | 10.5 | 18 | 9.5 | 4.7 | 3.72 | 27.56 | 46.86 | 5 |
| | B47 | 4 | 52 | 70 | 59 | 89 | 14.5 | 19 | 10.15 | 5.15 | 3.97 | 24.42 | 72.54 | 1 |
| | B48 | 4 | 39 | 59 | 72 | 105 | 14.5 | 14 | 9.65 | 5.36 | 3.86 | 25.18 | 30.22 | 5 |
| 40mM | C01 | 4 | 49 | 68 | 67 | 103 | 13.95 | 21 | 10 | 4.95 | 3.8 | 23.44 | 50.63 | 1 |
| | C02 | 3 | 43 | 59 | 38 | 68 | 14.05 | 21 | 9.25 | 4.69 | 3.71 | 17.7 | 77.01 | 2 |
| | C03 | 4 | 52 | 67 | 61 | 85 | 11.75 | 12 | 9.6 | 4.67 | 3.43 | 25.95 | 61.76 | 1 |
| | C04 | 4 | 40 | 55 | 50 | 76 | 11.5 | 17 | 10 | 5.5 | 4.18 | 25.96 | 68.52 | 1 |
| | C05 | 7 | 47 | 67 | 62 | 91 | 14.2 | 11 | 9.55 | 5.61 | 4.29 | 26.03 | 105.43 | 1 |
| | C06 | 7 | 45 | 70 | 66 | 92 | 11.95 | 12 | 9.85 | 4.84 | 3.68 | 25.34 | 44.6 | 5 |
| | C07 | 3 | 35 | 60 | 60 | 89 | 11.75 | 17 | 9.95 | 5.49 | 4.2 | 28.39 | 78.35 | 1 |
| | C08 | 4 | 46 | 62 | 67 | 106 | 14 | 11 | 9.95 | 5.48 | 4.09 | 23.9 | 55.22 | 1 |
| | C09 | 3 | 46 | 71 | 61 | 98 | 13.15 | 11 | 9.85 | 5.27 | 3.95 | 27.53 | 60.57 | 5 |
| | C10 | 4 | 55 | 76 | 68 | 97 | 16.75 | 12 | 9.75 | 5.09 | 4 | 25.64 | 57.43 | 1 |
| | C11 | 5 | 38 | 54 | 58 | 96 | 11 | 14 | 9.2 | 5.13 | 4.05 | 23.3 | 39.61 | 5 |
| | C12 | 6 | 44 | 59 | 67 | 98 | 13.6 | 18 | 9.85 | 5.56 | 3.97 | 26.7 | 82.24 | 1 |
| | C13 | 6 | 45 | 67 | 65 | 93 | 13.6 | 18 | 8.1 | 4.72 | 3.82 | 26.98 | 87.43 | 1 |
| | C14 | 3 | 43 | 68 | 39 | 74 | 14.95 | 17 | 9.4 | 4.63 | 3.49 | 17.77 | 52.79 | 2 |
| | C15 | 3 | 50 | 71 | 66 | 88 | 15.6 | 19 | 10 | 5.18 | 4.29 | 27.88 | 57.98 | 1 |
| | C16 | 7 | 52 | 69 | 55 | 80 | 12.5 | 17 | 9.6 | 5.14 | 3.98 | 27.98 | 76.93 | 1 |
| | C17 | 7 | 45 | 64 | 55 | 95 | 16.65 | 13 | 10.45 | 5.01 | 3.51 | 20.9 | 68.96 | 4 |
| | C18 | 5 | 53 | 78 | 66 | 91 | 12.6 | 19 | 9.65 | 4.87 | 3.99 | 26.66 | 72.78 | 1 |
| | C19 | 4 | 52 | 72 | 65 | 88 | 14.8 | 17 | 9.85 | 5.25 | 3.83 | 26.46 | 47.62 | 5 |
| | C20 | 4 | 46 | 71 | 47 | 83 | 13.7 | 13 | 9.3 | 4.79 | 3.51 | 21.5 | 54.19 | 5 |
| | C21 | 3 | 53 | 73 | 58 | 90 | 10.5 | 19 | 9 | 5.02 | 4.06 | 25.8 | 59.33 | 1 |

Values highlighted red indicate trait means significantly ($P<0.05$) below the control, while green indicate values significantly ($P<0.05$) above the control.Key: Trt= Treatment ID=Genotype Identifier, dtg days to germination, dff = days to first flower, d50f = days to 50% flowering, dfh = days to first harvest, d50mp = days to 50% mature pods, pdl = pod length, pdw = pod width, ln = number of locules, nsp = number of seeds per pod, psa = percent seed abortion, sdl = seed length, sdw = seed width, sdt = seed thickness, swgt = seed weight, yld = yield per plant

**S1 Table: Trait dataset and cluster grouping of individual $M_1$ cowpea plants across EMS treatments and control *(cont'd)***

| Trt | ID | *dtg* | *dff* | *d50f* | *dfh* | *d50mp* | *pdl* | *ln* | *sdl* | *sdw* | *sdt* | *swg* | yld | Cluster group |
|---|---|---|---|---|---|---|---|---|---|---|---|---|---|---|
| 40mM | C22 | 5 | 55 | 73 | 51 | 78 | 14.9 | 15 | 10 | 5.5 | 4.32 | 25.76 | 27.04 | 3 |
| | C23 | 5 | 43 | 64 | 55 | 81 | 12.05 | 18 | 9.6 | 4.79 | 3.84 | 21.7 | 42.96 | 1 |
| | C24 | 4 | 43 | 65 | 56 | 76 | 13.5 | 11 | 9.8 | 5.31 | 3.85 | 26.02 | 94.7 | 1 |
| | C25 | 4 | 40 | 59 | 62 | 102 | 12.5 | 12 | 9.4 | 4.81 | 3.63 | 23.51 | 55.96 | 5 |
| | C26 | 7 | 51 | 70 | 54 | 88 | 11.95 | 14 | 9.85 | 5.25 | 3.77 | 23.6 | 69.4 | 1 |
| | C27 | 5 | 48 | 69 | 67 | 95 | 6.95 | 19 | 9 | 4.58 | 5.59 | 23.01 | 48.33 | 1 |
| | C28 | 7 | 49 | 72 | 67 | 91 | 15.95 | 17 | 10 | 5.33 | 3.86 | 26.46 | 47.62 | 5 |
| | C29 | 5 | 39 | 62 | 56 | 92 | 15.4 | 13 | 10 | 4.98 | 3.81 | 25.45 | 42.76 | 5 |
| | C30 | 3 | 37 | 57 | 62 | 93 | 14.9 | 20 | 10 | 5.58 | 3.98 | 21.34 | 59.76 | 1 |
| | C31 | 5 | 42 | 59 | 50 | 83 | 12.75 | 18 | 10 | 4.88 | 3.57 | 27.56 | 48.24 | 5 |
| | C32 | 7 | 34 | 49 | 59 | 84 | 14.95 | 19 | 9.85 | 4.91 | 3.87 | 27.16 | 62.74 | 4 |
| | C33 | 3 | 52 | 77 | 50 | 76 | 12.25 | 19 | 9.35 | 5.33 | 4.21 | 24.58 | 63.91 | 1 |
| | C34 | 3 | 44 | 68 | 57 | 93 | 15.75 | 15 | 8.65 | 5.03 | 4.35 | 24.24 | 56.71 | 4 |
| | C35 | 5 | 48 | 70 | 58 | 94 | 15.45 | 18 | 9.85 | 4.88 | 4.01 | 23.82 | 60.04 | 1 |
| | C36 | 3 | 54 | 74 | 54 | 83 | 13.95 | 12 | 10.05 | 4.78 | 3.63 | 24.73 | 61.83 | 1 |
| | C37 | 4 | 49 | 67 | 51 | 76 | 13.9 | 17 | 9.8 | 5.28 | 4.08 | 23.83 | 31.45 | 1 |
| | C38 | 6 | 34 | 59 | 62 | 89 | 8.77 | 11 | 8.95 | 5.06 | 3.81 | 19.95 | 39.5 | 5 |
| | C39 | 6 | 37 | 57 | 52 | 92 | 15.5 | 19 | 10.1 | 4.98 | 3.91 | 22.71 | 27.02 | 5 |
| | C40 | 6 | 35 | 58 | 72 | 98 | 13.5 | 17 | 10 | 5.4 | 3.68 | 24.03 | 28.83 | 5 |
| | C41 | 5 | 55 | 80 | 64 | 89 | 14.5 | 13 | 10 | 4.83 | 3.71 | 26.36 | 90.96 | 1 |
| | C42 | 7 | 36 | 59 | 58 | 95 | 13.35 | 19 | 10 | 5.59 | 4.49 | 25.19 | 25.19 | 5 |
| | C43 | 5 | 35 | 53 | 71 | 95 | 14 | 15 | 10 | 5.55 | 4.45 | 27.28 | 90.01 | 1 |
| | C44 | 5 | 41 | 57 | 55 | 95 | 15.5 | 18 | 10 | 4.97 | 4.14 | 25.65 | 76.17 | 1 |
| | C45 | 3 | 40 | 59 | 54 | 84 | 15.75 | 12 | 11.05 | 5.07 | 3.8 | 27.77 | 83.32 | 1 |
| | C46 | 5 | 52 | 71 | 64 | 103 | 16.15 | 17 | 10.9 | 5.69 | 4.25 | 24.74 | 59.87 | 1 |
| | C47 | 4 | 40 | 57 | 63 | 87 | 13.95 | 11 | 10 | 5.58 | 4.09 | 23.3 | 37.51 | 3 |
| | C48 | 4 | 50 | 73 | 62 | 92 | 16.75 | 13 | 10 | 5.19 | 3.99 | 19.97 | 19.97 | 3 |
| | C49 | 7 | 39 | 55 | 59 | 80 | 14.75 | 8 | 10.4 | 5.29 | 3.92 | 22.82 | 60.26 | 3 |
| | C50 | 3 | 51 | 69 | 62 | 84 | 14.5 | 9 | 10.1 | 5.39 | 3.93 | 23.53 | 54.59 | 5 |
| | C51 | 4 | 53 | 70 | 62 | 87 | 12.5 | 13 | 9.9 | 5.3 | 3.96 | 19.66 | 21.63 | 3 |
| | C52 | 4 | 39 | 60 | 64 | 86 | 15.9 | 8 | 9.9 | 5.17 | 4.02 | 23.13 | 60.13 | 4 |
| | C53 | 6 | 39 | 63 | 62 | 85 | 14.75 | 9 | 9.75 | 5.22 | 3.9 | 19.28 | 75.19 | 4 |
| | C54 | 5 | 41 | 59 | 54 | 80 | 16.55 | 12 | 10.15 | 5.58 | 3.55 | 25.23 | 55.51 | 3 |
| | C55 | 7 | 53 | 70 | 51 | 89 | 14.5 | 11 | 9.85 | 4.76 | 3.69 | 25.06 | 52.62 | 5 |
| 80mM | D01 | 7 | 49 | 69 | 49 | 80 | 13.5 | 17 | 10 | 4.94 | 3.95 | 22.58 | 31.17 | 3 |
| | D02 | 6 | 50 | 72 | 54 | 92 | 10.75 | 10 | 10.7 | 5.71 | 4.28 | 24.18 | 42.55 | 1 |
| | D03 | 3 | 40 | 58 | 70 | 93 | 13.5 | 9 | 9.85 | 5.61 | 4.56 | 24.13 | 74.33 | 1 |
| | D04 | 5 | 55 | 73 | 72 | 100 | 12.5 | 13 | 9.35 | 5.34 | 3.81 | 18.82 | 76.2 | 1 |
| | D05 | 7 | 35 | 53 | 59 | 96 | 11.75 | 10 | 9.95 | 5.69 | 4.18 | 22.05 | 45.64 | 5 |
| | D06 | 4 | 34 | 59 | 55 | 93 | 15 | 9 | 10.3 | 5.68 | 4.16 | 20.79 | 50.32 | 3 |
| | D07 | 4 | 46 | 70 | 49 | 83 | 12.5 | 13 | 9.7 | 5.44 | 3.91 | 20.34 | 31.72 | 5 |
| | D08 | 4 | 48 | 69 | 61 | 81 | 15.25 | 13 | 10.55 | 6.11 | 4.94 | 26.24 | 43.3 | 3 |
| | D09 | 5 | 47 | 65 | 66 | 100 | 15 | 9 | 10.3 | 5.21 | 4.28 | 27.62 | 39.78 | 1 |
| | D10 | 4 | 53 | 78 | 58 | 97 | 13.75 | 10 | 9.85 | 5.04 | 4.38 | 24.53 | 35.57 | 3 |
| | D11 | 7 | 46 | 62 | 61 | 81 | 14 | 12 | 9.95 | 5.55 | 4.91 | 22.22 | 34 | 1 |
| | D12 | 3 | 43 | 68 | 59 | 95 | 15.75 | 9 | 10.05 | 5.58 | 4.7 | 22.36 | 60.82 | 6 |

Values highlighted red indicate trait means significantly ($P<0.05$) below the control, while green indicate values significantly ($P<0.05$) above the control.Key: Trt= Treatment ID=Genotype Identifier, dtg  days to germination, dff = days to first flower, d50f = days to 50% flowering, dfh = days to first harvest, d50mp = days to 50% mature pods, pel = peduncle length at first harvest, npp = number of pods per peduncle, nptm = number of pods per plant at maturity, nspt = number of seeds per plant, pdl = pod length, pdw = pod width, ln = number of locules, nsp = number of seeds per pod, psa = percent seed abortion, sdl = seed length, sdw = seed width, sdt = seed thickness, swgt = seed weight, yld = yield per plant

**S1 Table: Trait dataset and cluster grouping of individual $M_1$ cowpea plants across EMS treatments and control *(cont'd)***

| Trt | ID | *dtg* | *dff* | *d50f* | *dfh* | *d50mp* | *pdl* | *ln* | *sdl* | *sdw* | *sdt* | *swg* | yld | Cluster group |
|---|---|---|---|---|---|---|---|---|---|---|---|---|---|---|
| 80mM | D13 | 6 | 42 | 59 | 69 | 91 | 16.5 | 13 | 10.05 | 5.2 | 4.67 | 27 | 64.8 | 1 |
| 80mM | D14 | 6 | 52 | 74 | 45 | 81 | 12.75 | 12 | 9.4 | 5.37 | 4.46 | 24.66 | 54.24 | 1 |
| 80mM | D15 | 4 | 43 | 68 | 61 | 92 | 14.5 | 9 | 10.5 | 5.47 | 4 | 21.45 | 36.03 | 3 |
| 80mM | D16 | 5 | 48 | 68 | 65 | 92 | 13 | 13 | 9.9 | 5.02 | 3.95 | 22.74 | 68.23 | 1 |
| 80mM | D17 | 6 | 40 | 56 | 64 | 86 | 12.75 | 11 | 9.4 | 5.91 | 4.89 | 23.07 | 49.84 | 6 |
| 80mM | D18 | 3 | 44 | 68 | 64 | 101 | 11.45 | 7 | 9.95 | 4.94 | 3.97 | 25.55 | 35.77 | 1 |
| 80mM | D19 | 5 | 43 | 58 | 59 | 91 | 14.5 | 12 | 11 | 5.24 | 3.41 | 23.94 | 55.07 | 5 |
| 80mM | D20 | 4 | 55 | 75 | 64 | 103 | 11.25 | 13 | 10.75 | 5.41 | 4.21 | 22.04 | 65.9 | 1 |
| 80mM | D21 | 3 | 37 | 59 | 59 | 97 | 17.5 | 8 | 11 | 5.35 | 3.74 | 24.48 | 42.83 | 3 |
| 80mM | D22 | 7 | 47 | 62 | 63 | 92 | 14.5 | 14 | 9.35 | 4.73 | 3.74 | 20.59 | 40.77 | 4 |
| 80mM | D23 | 7 | 37 | 55 | 57 | 78 | 16 | 13 | 9.35 | 5.21 | 3.94 | 22.7 | 25.88 | 3 |
| 80mM | D24 | 4 | 40 | 62 | 72 | 109 | 14.25 | 8 | 9.15 | 4.95 | 4.07 | 17.35 | 58.65 | 4 |
| 80mM | D25 | 3 | 45 | 68 | 68 | 90 | 15 | 9 | 11 | 5.62 | 4.15 | 24.44 | 102.66 | 1 |
| 80mM | D26 | 4 | 46 | 63 | 71 | 110 | 17.1 | 13 | 8.95 | 5.02 | 3.86 | 22.7 | 22.7 | 3 |
| 80mM | D27 | 4 | 44 | 62 | 61 | 86 | 16.25 | 8 | 10 | 5.31 | 4.04 | 21.91 | 42.73 | 4 |
| 80mM | D28 | 6 | 43 | 66 | 60 | 92 | 17.1 | 9 | 10.45 | 5.48 | 4.28 | 22.32 | 75 | 4 |
| 80mM | D29 | 7 | 39 | 62 | 58 | 82 | 12.75 | 12 | 7.95 | 4.28 | 3.45 | 21.58 | 69.93 | 4 |
| 80mM | D30 | 4 | 35 | 60 | 56 | 84 | 12.75 | 11 | 10.25 | 5.02 | 3.79 | 25.06 | 36.08 | 5 |
| 80mM | D31 | 4 | 35 | 55 | 54 | 87 | 14.9 | 17 | 10.75 | 5.17 | 4.02 | 21.98 | 24.17 | 5 |
| 80mM | D32 | 4 | 44 | 62 | 68 | 96 | 14.15 | 11 | 10.2 | 6.05 | 4.82 | 22.16 | 37.23 | 1 |
| 80mM | D33 | 4 | 37 | 60 | 48 | 85 | 11.25 | 7 | 10.3 | 4.81 | 3.92 | 25.86 | 46.54 | 5 |
| 80mM | D34 | 4 | 45 | 66 | 52 | 83 | 12.25 | 12 | 10.25 | 4.97 | 3.46 | 21.21 | 23.75 | 3 |
| 80mM | D35 | 7 | 41 | 66 | 58 | 81 | 16.75 | 13 | 9.3 | 5.09 | 4.06 | 21.79 | 35.96 | 3 |
| 80mM | D36 | 3 | 53 | 71 | 49 | 73 | 14 | 8 | 11 | 5.64 | 4.37 | 28.29 | 74.69 | 6 |
| 80mM | D37 | 4 | 43 | 61 | 59 | 80 | 15.25 | 9 | 10 | 5.38 | 4.08 | 25.05 | 50.86 | 5 |
| 80mM | D38 | 5 | 42 | 57 | 62 | 88 | 15.5 | 10 | 9.3 | 5.14 | 3.5 | 17.49 | 44.24 | 4 |
| 80mM | D39 | 4 | 40 | 56 | 52 | 90 | 15.75 | 9 | 9.4 | 5.3 | 4.06 | 19.27 | 48.57 | 3 |
| 80mM | D40 | 4 | 44 | 69 | 54 | 77 | 9.95 | 13 | 8.55 | 5.18 | 3.96 | 19.42 | 55.55 | 2 |
| 80mM | D41 | 7 | 45 | 63 | 59 | 81 | 15 | 12 | 9 | 4.79 | 3.75 | 19.76 | 80.03 | 4 |
| 80mM | D42 | 3 | 55 | 80 | 66 | 96 | 9.95 | 11 | 9.9 | 4.72 | 3.72 | 22.02 | 29.06 | 3 |
| 80mM | D43 | 3 | 54 | 73 | 48 | 69 | 11.7 | 17 | 10.5 | 4.9 | 3.67 | 21.98 | 42.19 | 1 |
| 80mM | D44 | 6 | 48 | 65 | 52 | 86 | 9.5 | 12 | 11.6 | 6.03 | 4.46 | 24.35 | 64.29 | 1 |
| 80mM | D45 | 4 | 47 | 68 | 60 | 92 | 11.5 | 9 | 9.8 | 5.4 | 4.22 | 23.58 | 106.09 | 1 |
| 80mM | D46 | 3 | 44 | 62 | 57 | 78 | 12.5 | 13 | 9.8 | 5.05 | 4.15 | 21.53 | 45.21 | 3 |
| 80mM | D47 | 3 | 38 | 59 | 58 | 78 | 13 | 13 | 9.85 | 5.08 | 3.99 | 18.45 | 18.45 | 3 |
| 80mM | D48 | 5 | 47 | 65 | 52 | 84 | 13.65 | 8 | 10 | 5.3 | 4.24 | 22.52 | 85.13 | 1 |
| 80mM | D49 | 7 | 52 | 72 | 62 | 83 | 11.2 | 9 | 10.45 | 5.48 | 4.03 | 22.93 | 79.1 | 1 |
| 80mM | D50 | 3 | 42 | 60 | 52 | 92 | 13.1 | 12 | 9.65 | 4.82 | 3.7 | 22.5 | 43.19 | 5 |
| 80mM | D51 | 7 | 33 | 56 | 50 | 83 | 8.45 | 11 | 9.55 | 4.76 | 3.84 | 22.62 | 64.7 | 5 |
| 80mM | D52 | 5 | 55 | 77 | 56 | 84 | 14.2 | 17 | 10.4 | 4.87 | 3.87 | 19.54 | 22.48 | 3 |
| 80mM | D53 | 7 | 43 | 62 | 59 | 99 | 11.4 | 12 | 9.8 | 5.51 | 4.38 | 22.83 | 55.25 | 1 |
| 80mM | D54 | 5 | 47 | 71 | 60 | 85 | 11.5 | 9 | 9.8 | 5.44 | 4.1 | 20.24 | 28.33 | 3 |
| 80mM | D55 | 4 | 37 | 62 | 43 | 74 | 14.25 | 13 | 9.05 | 4.85 | 3.85 | 20.31 | 35.34 | 2 |
| 80mM | D56 | 5 | 53 | 69 | 57 | 94 | 13.5 | 23 | 10.15 | 5.03 | 3.78 | 28.68 | 111.85 | 1 |

Values highlighted red indicate trait means significantly ($P<0.05$) below the control, while green indicate values significantly ($P<0.05$) above the control.Key: Trt= Treatment ID=Genotype Identifier, dtg = days to germination, dff = days to first flower, d50f = days to 50% flowering, dfh = days to first harvest, d50mp = days to 50% mature pods, pdl = pod length, pdw = pod width, ln = number of locules, nsp = number of seeds per pod, psa = percent seed abortion, sdl = seed length, sdw = seed width, sdt = seed thickness, swgt = seed weight, yld = yield per plant
