## Supplementary material for "Effect of Ethyl Methane Sulfonate Mutagenesis on Phenological, Yield-Related and Yield Traits in Cowpea *(Vigna unguiculata* (L.) Walp)": S2 Table

### S1 Table: PCA loadings and variance explained for traits in $M_1$ cowpea

| Traits | PC1 | PC2 | PC3 | PC4 | PC5 | PC6 | PC7 | PC8 | Uniqueness |
|---|---|---|---|---|---|---|---|---|---|
| sdl | 0.73 | | | | | | | | 0.18 |
| sdw | 0.70 | | 0.50 | | | | | | 0.11 |
| dfh | 0.68 | | | 0.50 | | | | -0.32 | 0.14 |
| dff | 0.61 | | -0.64 | | | | | | 0.10 |
| d50mp | 0.59 | | | 0.51 | | | | -0.40 | 0.13 |
| d50f | 0.57 | | -0.68 | | | | | | 0.09 |
| swg | 0.53 | | | | 0.35 | | | 0.43 | 0.36 |
| sdt | 0.38 | | 0.42 | | | 0.52 | -0.38 | | 0.24 |
| nspt | | 0.96 | | | | | | | 0.02 |
| yld | | 0.95 | | | | | | | 0.02 |
| nsp | | 0.75 | | 0.36 | -0.38 | | -0.32 | | 0.03 |
| nptm | | 0.32 | | -0.66 | | | 0.53 | | 0.04 |
| pdl | | | 0.42 | | | | | | 0.77 |
| pdw | | | | | 0.54 | -0.35 | | 0.32 | 0.40 |
| ln | | | | | 0.47 | 0.63 | | | 0.22 |
| dtg | | | | | 0.46 | -0.33 | | | 0.51 |
| psa | | | | | -0.42 | | | 0.32 | 0.52 |
| npp | | | | | | 0.32 | 0.41 | 0.43 | 0.43 |
| **Eigenvalue** | **3.18** | **2.71** | **1.70** | **1.51** | **1.34** | **1.17** | **1.07** | **1.03** | |
| **Proportion %** | **17.70** | **15.10** | **9.40** | **8.40** | **7.50** | **6.50** | **6.00** | **5.70** | |
| **Cumulative %** | **17.70** | **32.70** | **42.20** | **50.60** | **58.00** | **64.50** | **70.50** | **76.20** | |

PC= Principal component, sdl = seed length, sdw = seed width, dfh = days to first harvest, dff = days to first flower, , d50f = days to 50% flowering, , d50mp = days to 50% mature pods, swgt = seed weight, sdt = seed thickness, nspt = number of seeds per plant npp, yld = yield per plant, nsp = number of seeds per pod, nptm = number of pods per plant at maturity, pdl = pod length, pdw = pod width, ln = number of locules dtg = days to germination, psa = percent seed abortion, npp= number of pods per peduncle
